# Connectome simulations identify a central pattern generator circuit for fly walking

**DOI:** 10.1101/2025.09.12.675944

**Authors:** Sarah M. Pugliese, Grant M. Chou, Elliott T.T. Abe, Denis Turcu, Jackson K. Lancaster, John C. Tuthill, Bingni W. Brunton

## Abstract

Animal locomotion relies on rhythmic body movements driven by central pattern generators (CPGs): neural circuits that produce oscillating output without oscillating input. However, the circuit structure of a CPG for walking is not known in any animal. To identify the cells and synapses that underlie rhythmic leg movement in walking flies, we developed dynamic simulations of the *Drosophila* ventral nerve cord (VNC) connectomes. A computational activation screen of descending neurons from the central brain identified DNg100—a known command neuron for walking—as the top driver of rhythmic leg motor activity. Simulated network pruning isolated a minimal rhythm-generating circuit consisting of one inhibitory and two excitatory interneurons; this three-neuron circuit was necessary and sufficient for motor rhythms across all six legs and in four connectome datasets. Simulations also predicted that a separate descending pathway (DNb08) drives rhythmic leg movements, which we confirmed experimentally using optogenetics in behaving flies. Our results reveal the cellular identity and synaptic structure of a putative CPG circuit for walking and other rhythmic leg movements in flies.

---

Most animals move through the world using rhythmic movements, such as undulation of the body to swim and slither, or cyclic movement of legs and wings to walk or fly. Early studies of vertebrate locomotion proposed that motor rhythms are generated by sensory feedback from proprioceptors, which reflexively trigger the next phase of the locomotor cycle [1]. However, subsequent work on cats [2], crayfish [3], and locusts [4] demonstrated that motor rhythms persisted even after transection of sensory nerves. These results suggested that the central nervous system can generate rhythmic activity patterns on its own, without proprioceptive feedback or other rhythmic inputs. Such rhythm-generating circuits became known as *central pattern generators* (CPGs). CPGs have been studied intensively in both vertebrates and invertebrates, where they are crucial in locomotor rhythms such as walking, flying, and swimming, and non-locomotor rhythms such as breathing, feeding, scratching, circulation, and singing [5, 6, 7].

CPG circuits have been described to generate rhythms using multiple cellular and circuit mechanisms. For example, in the crustacean stomatogastric ganglion (STG, [8]) as well as the leech heart and swimming CPGs [9, 10], neurons possess specialized cellular properties that support rhythm generation, such as intrinsic bursting, plateau potentials, and post-inhibitory rebound. The preBö tzinger complex, which controls breathing in vertebrates, contains neurons that intrinsically burst, even when isolated from the rest of the network [11]. However, specific patterns of synaptic connectivity between neurons, such as mutual inhibition, can also contribute to rhythm generation in these circuits [12]. For example, the swimming CPG of the nudibranch *Tritonia* is a network oscillator whose cells lack intrinsic bursting properties [13]. Overall, studies of diverse CPG circuits suggest that rhythm generation is often an emergent property of the network as a whole, rather than being reliant on one specific mechanism [14, 15].

Despite considerable evidence that CPGs produce the walking rhythm in vertebrates and invertebrates, the specific implementation of a CPG circuit for walking is not known for any animal [5, 7]. Lacking knowledge of the anatomy, connectivity, and electrophysiological properties of CPG neurons, most computational models for animal walking rely on the presence of intrinsic bursting neurons as components of balanced excitation-inhibition networks (e.g., [16, 17]), or abstract all neural mechanisms to replace the CPG circuit with an oscillator equation (e.g., the Kuramoto oscillator [18]). Much theoretical work has focused on how CPGs interact within and across limbs [19, 20, 21, 22] and with the body [23, 24, 25] to produce coordinated locomotion, without explicitly modeling the underlying pattern-generating mechanisms.

Here, we seek to identify the cellular components of CPG circuits that underlie walking in the fruit fly, *Drosophila*, using a computational modeling approach enabled by comprehensive datasets of neural connectivity. Flies are agile and robust walkers, and the adult fly is the only limbed animal whose nervous system is almost completely mapped at synaptic resolution using electron microscopy reconstruction of cells and synapses [26, 27, 28, 29, 30, 31]. Recent connectome datasets that include the fly ventral nerve cord (VNC, [28, 29, 30, 31]), which is analogous to the vertebrate spinal cord (**Fig. 1a**), provide a unique opportunity to identify the core cell types and synaptic connections that comprise fly CPGs. Beyond synaptic connectivity maps, published VNC connectome datasets have been extensively annotated [32, 33, 34] and include predictions of neurotransmitter identity and muscle targets of leg motor neurons.

**Figure 1.**
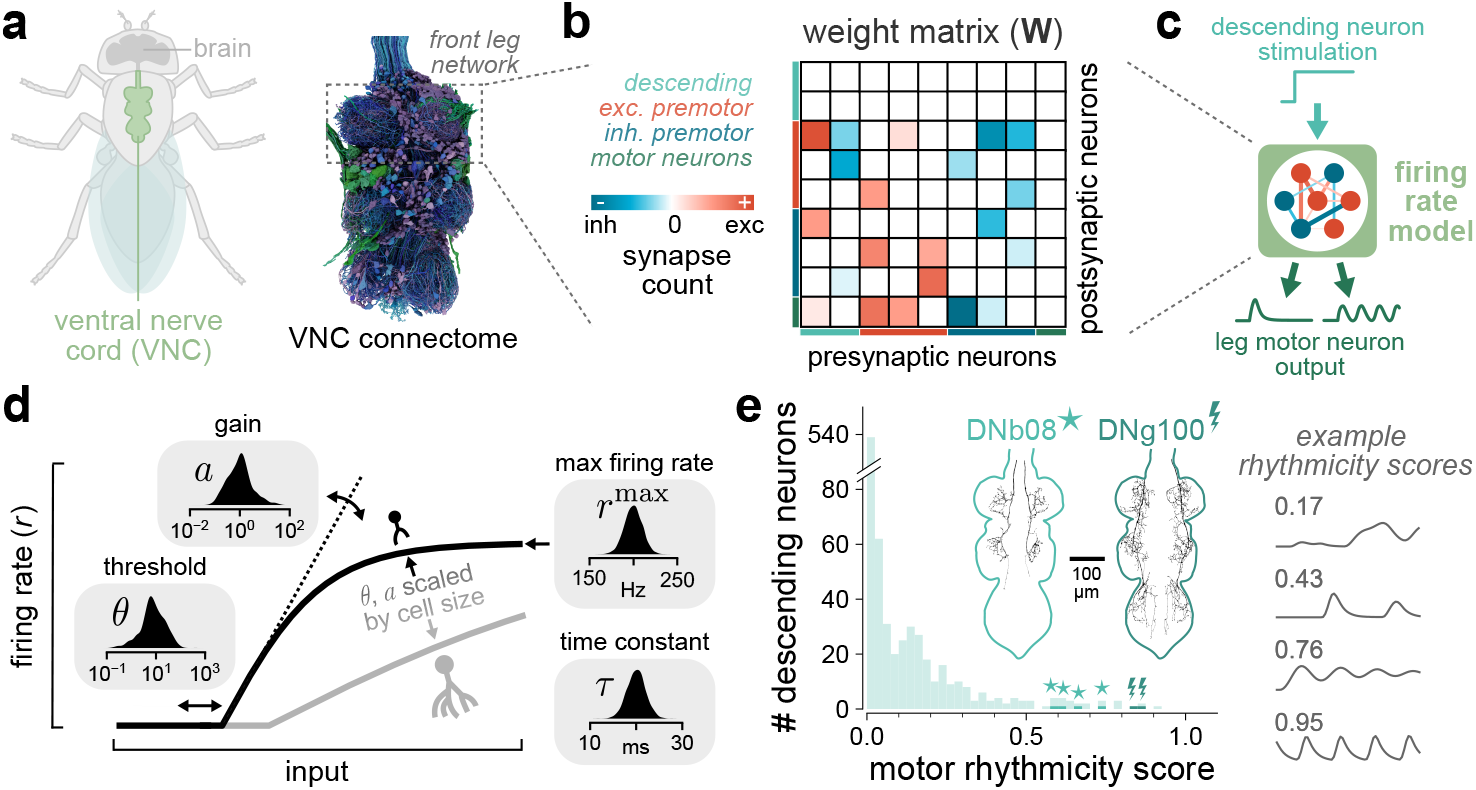
Direct numerical simulations of fly ventral nerve cord (VNC) connectomes identify descending pathways for rhythmic leg motor control. **a**, The *Drosophila* VNC, which is analogous to the vertebrate spinal cord. We constructed network models based on the connectivity of cells in the front leg neuropil from the available adult ventral nerve cord connectomes (dashed box). **b**, Schematized synaptic weight matrix **W**, where red and blue indicate excitatory and inhibitory connections, respectively. **c**, Schematic of model and descending neurons (DNs) activation screen. Each excitatory DN was stimulated with tonic input, which then activated a rate model of the VNC connectome (**Equation 1**). We assessed the rhythmicity of leg motor neuron rates. **d**, Neurons in the model had four biophysical parameters and a rectified tanh activation function. Insets show the distributions from which these parameters for each neuron were drawn randomly at every simulation replicate. Gain *a* and threshold *θ* were scaled by the size of each neuron (see **Methods**). **e**, A computational screen of DNs identified cells that produce rhythmic activity in leg motor neurons. Shown at right are example motor neuron activity traces and their rhythmicity scores. Each DN’s score was an average of 128 replicates. Two top-scoring DN types are highlighted (DNb08 as stars, DNg100 as bolts, with their anatomy from MANC shown as insets).

Based on analyses of small nervous systems mapped using connectomics (e.g., the *C. elegans* nematode) and electrophysiology (e.g., the crustacean STG), it has been argued that computational models of neural dynamics are not sufficiently constrained by connectivity alone [35, 36]. Existing fly connectome datasets lack important biophysical and molecular parameters, such as the expression of ion channels, receptors, neuromodulators, and gap junctions. Despite these gaps, several recent studies have developed connectome-constrained simulations of circuits within the fly central brain. One study directly modeled neural activity using a spiking network with fixed parameters [37], while others have leveraged neural recording data and task constraints to infer unknown parameters with machine learning [38, 39, 40]. Here, we developed direct numerical simulations of fly VNC connectomes to identify descending pathways that produce rhythmic leg movement during sustained activation from the brain, and corroborated these predictions with optogenetics experiments in behaving flies. We then developed a computational pruning screen to isolate a minimal circuit of three VNC interneurons that is necessary and sufficient to generate rhythmic leg motor activity. We propose that this three-neuron circuit constitutes a core CPG for walking and other rhythmic leg movements in flies.

## Connectome simulations of motor activity from descending input

To identify rhythm-generating circuits for fly walking, we developed rate models of neural activation based on the four published adult fly connectome datasets that include the VNC [28, 29, 30, 31]. We first focused our analysis on the connectivity of neurons controlling the front legs in the male adult nerve cord (MANC) dataset [28]. This simulated network consisted of 4,604 neurons, including 1,318 descending neurons (DNs), 144 leg motor neurons, and 3,142 premotor neurons within the VNC, which we defined as all non-descending neurons that synapse onto leg motor neurons [32]. The 3,817,772 synapses among these neurons formed elements of a weight matrix **W**, for which the entries **w**_*ij*_ are the signed synapse counts from presynaptic neuron *j* to postsynaptic neuron *i* (**Fig. 1b**). To assign synaptic weights as positive or negative, we used the predicted neurotransmitter identity of neuron *j*, such that excitatory (cholinergic) cells had positive weights and inhibitory (GABAergic, glutamatergic) cells had negative weights. We chose a rate model in part because many neurons in the insect VNC, including premotor neurons active during walking, are nonspiking [41, 42, 43, 44].

We modeled the rates of neurons in the network **r**_*i*_(*t*) as a system of nonlinear ordinary differential equations, given their synaptic and external inputs. Rates evolved according to an equation of the form 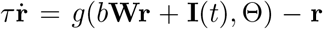, where *g*(·) is an input nonlinearity specified by cellular parameters Θ: *a, θ*, and *r*^max^, described below (**Fig. 1d**, full form in **Eq. 1**). The input to neuron *i* was a sum of rates {**r**_*j*_(*t*)} weighted by signed synapse counts **w**_*ij*_, and any external input **I**_*i*_(*t*) to the neuron. A constant *b* scaled the synaptic inputs relative to the external input and had the same value for all neurons. This summed input was then passed through a nonlinearity *g*(·) such that rates below threshold *θ* were zero and gradually increased with gain *a* until saturation at a maximum value *r*^max^. The rate of each neuron decayed to zero with time constant *τ* . We normalized the gain and threshold parameters to account for the fact that larger neurons have proportionally lower input resistance and are therefore less excitable [45]. We based this size normalization on experimental data showing that postsynaptic potential amplitudes scale linearly with the number of synapses per unit area in *Drosophila* neurons [46].

In each replicate of the simulations, for each neuron *i*, we randomly drew its biophysical parameters {*a*_*i*_, *θ*_*i*_, *τ*_*i*_, 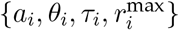} from distributions based on published physiological measurements of fly neurons [44, 47]; values were also consistent with previous fly network models [37, 48]. Rather than fine-tuning parameters, we simulated large numbers of replicates with independently drawn biophysical parameters, so that consistent dynamics across replicates indicate that the functions arise from synaptic connectivity structure rather than precise knowledge of any of the 18,416 parameters in the network. Unlike past studies that exhaustively varied cellular and synaptic parameters to characterize the space of configurations supporting a given network output [49], we fixed connectivity to the exact connectome dataset weights and varied biophysical parameters across simulation replicates to demonstrate robustness. Note that no neuron in our models had intrinsic bursting or other longer-timescale membrane properties—all dynamics emerged from the rate model rather than relying on specialized cellular properties. Further details on the construction and numerical implementation of simulations are elaborated in **Methods**.

To identify which descending neurons drive rhythmic leg movement, we conducted a computational activation screen of all 933 excitatory DNs (i.e., DNs predicted to release acetylcholine). Because the model had no spontaneous activity, activating inhibitory DNs alone could not recruit downstream neurons, so they were excluded from the screen. We activated each DN with a step input and evaluated average output rhythmicity of leg motor neurons over 128 random simulation replicates (**Fig. 1c**). A small fraction of DNs consistently produced highly rhythmic activity across replicates, suggesting they are high-confidence candidates for descending inputs to a rhythm-generating circuit (**Fig. 1e**).

To quantify the rhythmicity of simulated motor neuron activity, we developed a *motor rhythmicity score* that was calculated from the autocorrelations of active motor neurons, averaged over all replicates (see **Methods**). The motor rhythmicity score is a number between 0 and 1, where 0 is not rhythmic and 1 corresponds to perfectly periodic oscillations (see example traces in **Fig. 1e**). Out of the 933 DNs activated, 74 (7.9%) produced unstable simulations in all replicates so could not be scored (see **Methods**). Of the 859 remaining, 332 (38.6%, or 35.6% of the total) had an average score of 0, and 207 more scored below 0.025 on average (24.1%, or 22.2% of total). Only 29 DNs (3.4%, or 3.1% of total) had an average score greater than 0.5 (**Supplementary Table 1**). DN activation screen results were qualitatively similar across a wide range of model parameters and configurations, including activating DNs of the same cell type (rather than one-by-one, **Extended Data Fig. 1c**), combinatorially activating multiple DNs (**Extended Data Fig. 1e** and **Supplementary Table 2**), and increasing the variance of all four biophysical parameter distributions (**Extended Data Fig. 2c**).

Together, these results demonstrate that connectome simulations can identify candidate descending pathways for rhythmic leg movements. We note that this computational screen works as a discovery tool, analogous to a forward genetic screen, to identify high-confidence candidate DNs for further investigation. Like a forward genetic screen, it likely produced false negatives; therefore, we do not interpret low scores as evidence against a DN’s role in rhythm generation. In subsequent analyses, we focus on two descending neuron types, DNg100 and DNb08, because every DN of these two types produced among the highest rhythmicity scores across all simulation replicates.

### DNg100 activation drives rhythmic leg motor neuron activity and controls stepping frequency

DNg100 is the only cell type in *Drosophila* known to function as descending command neurons for walking; optogenetic stimulation of DNg100 neurons initiates forward walking, even in headless flies ([50], where DNg100 is referred to as BDN2). To investigate how DNg100 activation propagates through the VNC to produce rhythmic motor output, we simulated DNg100 activation in each of the four published VNC connectome datasets: the male (MANC, [28]) and female (FANC, [29]) adult nerve cord networks, the male central nervous system (mCNS, [31]), and the female brain and nerve cord (BANC, [30]). For each dataset, we simulated 1024 replicates where a single sustained input was applied to the DNg100 neuron innervating the left side of the VNC. Across all four connectomes, DNg100 reliably drove rhythmic activity in front leg motor neurons (**Fig. 2a–d**), with comparable rhythmicity scores despite some variation in the number of recruited motor neurons (**Fig. 2e**). The consistency of these results across four independent datasets from two sexes suggests that the VNC network downstream of DNg100 contains a conserved CPG circuit.

**Figure 2.**
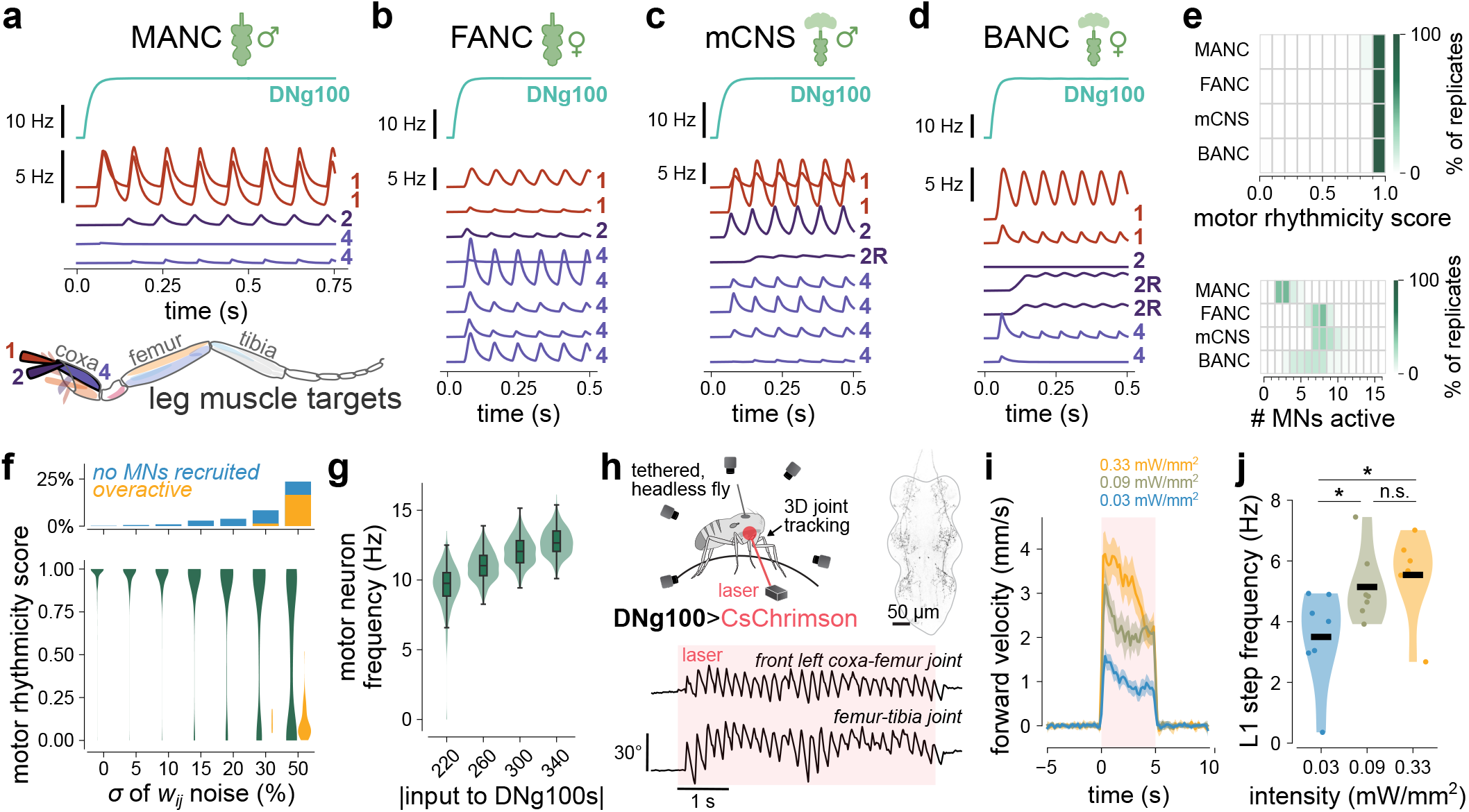
DNg100 produces rhythmic motor neuron activity in network simulation of four connectome datasets and in headless flies. **a**, Example activity of leg motor neurons (MNs) in MANC network with DNg100 activation. Numbers correspond to muscles innervated by each MN. In this plot and all other example trace plots, the simulation begins as a quiescent network (rates at *t* = 0 s are 0 Hz). **b-d**, Same as **a** for the FANC, mCNS, and BANC connectomes, respectively. **e**, *Top*, DNg100 activation consistently produced rhythmic motor activity in all four connectome datasets. *Bottom*, DNg100 typically recruited 2–10 leg MNs. For each dataset, *n*=1024 replicates. **f**, Motor rhythms were robust to noise perturbations of weights in the connectivity matrix up to 10% (*n*=512 replicates per noise condition); increasing noise led to more replicates with no MN recruitment or unstable dynamics (*top*). **g**, Increasing amplitude of input to DNg100 increased motor neuron oscillation frequency in MANC network (*n*=512 replicates per condition). Simulations with no active motor neurons were omitted. Green shaded regions are distributions of frequencies, and black bars indicate medians and quartiles. **h**, Example coxa-femur and femur-tibia joint kinematics of front left leg during optogenetic activation of DNg100 neurons in a tethered headless fly walking on a spherical treadmill (see also **Supplementary Video 1**). Top right shows confocal image of VNC expression (DNg100*>*CsChrimson, [50]). **i**, At higher optogenetic stimulus intensities, flies walked with higher forward velocity (mean ±95% c.i., *n*=7 flies, 8 trials per fly). **j**, Stepping frequency of the left front leg increased with laser intensity (*p* = 0.032, 0.352, and 0.042 for paired *t*-tests comparing low-medium, medium-high, and low-high laser intensities, respectively).

To determine the extent to which leg motor rhythms depend on the precise synaptic weights in the connectomes, we tested the robustness of DNg100-driven rhythms by systematically adding noise to the weight matrix. Specifically, we altered **W** from the MANC dataset by adding noise perturbations *η*_*ij*_ to each entry **w**_*ij*_. The magnitude of perturbations *η*_*ij*_ was drawn from a normal distribution with standard deviation proportional to **w**_*ij*_ (so that *η*_*ij*_ ∼ 𝒩 (0, *σ***w**_*ij*_), where *σ* varied from 0 to 50%), truncated so that the perturbation cannot change the sign of **w**_*ij*_, thus preserving the structure of the connectivity matrix. Motor rhythms were robust up to 10% added noise (**Fig. 2f**), which is comparable to the estimated uncertainty in synapse counts arising from both biological variability across individuals and errors in automated synapse detection [26, 51]. That rhythms persist under perturbations at the scale of connectome measurement error suggests that the rhythm-generating capacity of the DNg100 circuit is a robust property of its overall connectivity structure, and does not depend on precise synapse counts.

The oscillations produced in leg motor neurons downstream of DNg100 roughly matched the stepping frequency of real walking flies (∼ 7–15 Hz, [52, 53]) using biologically-plausible time constants (*τ*) with a mean of 20 ms (see **Methods** for a mathematical relationship between frequency and *τ* in the linearized system). Notably, increasing the magnitude of DNg100 stimulation increased the frequency of motor neuron oscillations in simulation (**Fig. 2g**), generating a testable prediction: stronger descending drive from DNg100 should produce faster stepping. Although previous work had shown that optogenetically activating DNg100 neurons is sufficient to drive walking in headless flies ([50]), the effect of stimulus intensity on stepping frequency had not been explored. To test this prediction, we expressed CsChrimson in DNg100 neurons and then optogenetically activated them in decapitated flies on a spherical treadmill (**Fig. 2h, Extended Data Fig. 3, Supplementary Video 1**). Increasing laser intensity led to an increase in both forward velocity and stepping frequency (**Fig. 2i–j**). This experiment confirmed predictions from our connectome simulations and suggests that the oscillation frequency of the CPG for walking is controlled by the strength of descending input from DNg100.

### A core CPG circuit of three interneurons

Having established that DNg100 drives rhythmic leg motor activity through the VNC network, we next sought to identify the specific cells and synapses responsible for transforming this sustained descending command into rhythmic motor output. The circuitry linking DNg100 to leg motor neurons is extensive and highly recurrent (**Fig. 3a**), and we anticipated that this recurrence is crucial because feedforward circuits alone cannot produce oscillating outputs from tonic inputs. Identifying the rhythm-generating subcircuit within this densely connected network would be intractable using conventional connectome analyses that trace feedforward pathways, since such approaches do not account for the dynamics that arise from recurrent connectivity.

**Figure 3.**
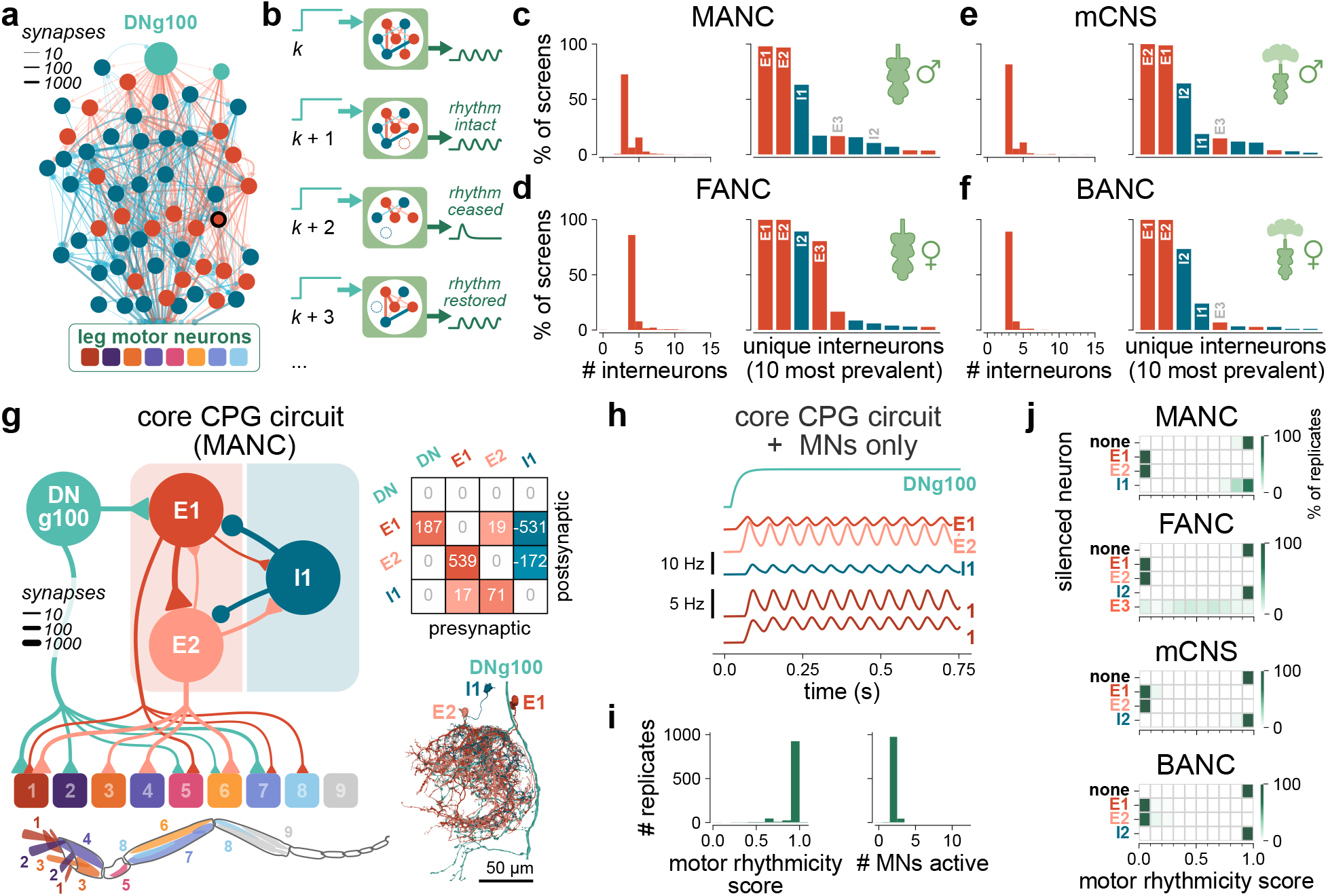
A computational sufficiency screen isolates a core CPG circuit downstream of DNg100. **a**, Premotor network downstream of DNg100 in MANC. Shown are all neurons with ≥10 synapses from left DNg100 or to any left front leg motor neuron. Red cells are putative excitatory (cholinergic), blue cells are inhibitory (GABAergic or glutamatergic). E1 is outlined in black. **b**, Schematic of iterative pruning procedure under DN input (teal). At each iteration, we removed one cell (dashed outlines) stochastically and evaluated the motor rhythmicity. This procedure terminated at a minimal circuit when no further neurons can be pruned without the rhythmicity score dropping below 0.5 (evaluated on MN output, green traces). **c**, Minimal circuits identified by pruning screens (*n*=1024 replicates, left DNg100, MANC). *Left*: Number of interneurons remaining in minimal circuits; 5 screens that did not converge to 15 or fewer interneurons are excluded. *Right*: Percentage of screens in which minimal circuits contained each unique interneuron (see **Supplementary Table 3** for full list of cell identities). **d–f**, Same as **c**, but for FANC (1 outlier of 18 interneurons not shown), mCNS, and BANC, respectively. See **Supplementary Tables 4–6** for full list of cell identities. **g**, The core CPG circuit in MANC and its connectivity to motor neurons (shown are connections of at least 10 synapses). Triangles and circles are excitatory and inhibitory connections, respectively. Rounded boxes depict modules of motor neurons grouped by leg muscle innervation [32], illustrated on the leg schematic below. Synapse counts are summed across MNs within each module. Top inset shows the core CPG weight matrix. Bottom inset shows anatomy of the core CPG circuit neurons (MANC) in the front left leg neuropil viewed laterally. **h**, DNg100 activation of the isolated three-neuron CPG circuit produced rhythmic activity in interneurons (E1, E2, I1; *top*) and leg motor neurons (*bottom*). **i**, The circuit in **g** consistently produced motor rhythms (*n*=1024 replicates). *Left*: distribution of motor rhythmicity scores. *Right*: number of active motor neurons per replicate. **j**, In the full network simulations, individually silencing E1 and E2 abolished leg motor rhythms, but silencing I1 (MANC), I2 (FANC, mCNS, BANC), and E3 (FANC) did not.

We therefore developed a *computational sufficiency screen* to isolate the minimal subset of neurons that form a circuit capable of generating rhythmic motor output. Starting from the full network simulations described above (**Fig. 2**), our screen proceeded by iterative pruning. At each iteration, we stochastically silenced one interneuron, selected with a probability inversely proportional to its activity. If leg motor oscillations persisted (rhythmicity score *>*0.5), that cell was pruned from the network . However, if the oscillations ceased (rhythmicity *<*0.5), the cell was restored (**Fig. 3b**, see **Methods** for additional details). Each replicate of the screen terminated when none of the remaining cells could be pruned without disrupting leg motor neuron rhythms. The goal of the screen was to prune the network to a minimal, core subcircuit sufficient to generate leg motor rhythms in response to sustained DNg100 activation.

In MANC, independent computational sufficiency screens repeatedly converged to a minimal circuit of just three neurons, two excitatory and one inhibitory (**Fig. 3c**). Over 60 percent (636/1024) of pruning screens converged to this same circuit. These three cells, which we refer to as “E1” (IN17A001), “E2” (INXXX466), and “I1” (IN16B036) for clarity, are interneurons local to the front left leg neuropil. In FANC, screens converged to a four-interneuron circuit (70.4% of 1024 replicates) that retained E1 and E2 but identified a different inhibitory neuron, “I2” (IN19A007), and one additional excitatory interneuron, “E3” (IN19B012, **Fig. 3d**). In the two full CNS connectomes, pruning screens converged to circuits containing either I1 or I2 alongside E1 and E2 (**Fig. 3e,f**). Despite the stochastic nature of the pruning procedure and differences across datasets, the minimal circuits nearly always included both E1 and E2, and either I1 or I2 (3521/4096 screens, or 86.0%). **Supplementary Tables 3–6** contain a detailed list of the distinct circuits and cells identified by all of the pruning screens.

In the MANC minimal circuit, the three neurons E1, E2, and I1 are connected all-to-all, but with particularly strong connections from E1→ E2, E2 →I1, I1 →E1, and I1 →E2 (**Fig. 3g**). Only E1 receives direct synaptic input from DNg100. Thus, the predominant mechanism for rhythm generation is that DNg100 drives E1 to excite E2, amplifying the excitation, and E2 then recruits I1 to inhibit both excitatory neurons after some delay (corresponding to the time-constant parameter *τ* in our model). Since I1 receives no direct input from DNg100, the inhibition onto E1 is eventually released, allowing E1 to reactivate and restart the cycle. This pattern repeats cyclically as long as DNg100 provides sustained excitation. Consistent with this mechanism, a reduced model of the three-neuron circuit that retains only the E1 →E2, E2 →I1, and I1 →E1 connections still produced oscillatory activity (**Extended Data Fig. 4b**). The pattern of connectivity of E1 and E2 with the neuron I2 is qualitatively similar to the connectivity with I1 (see **Extended Data Fig. 4g** for synaptic weights of these neurons in all 4 connectomes), consistent with the partial degeneracy of I1 and I2 observed across pruning screens. Finally, because both E1 and E2 connect to multiple motor neurons innervating muscles throughout the leg, their differential activation patterns drive motor neurons at consistent phase offsets relative to each other (**Fig. 3g,h**).

The recurrent E-E-I connectivity motif is common in the VNC connectome: we found 21,544 instances of this motif in the MANC front leg network, far exceeding the 307 ±29 (mean± std) expected from random shuffles of the weight matrix. What distinguishes the E1-E2-I1 circuit from the vast majority of these other instances is the exceptional strength of its recurrent connections (**Extended Data Fig. 4e**). Among the cumulative input synapses to E1, E2, and I1, 9.28% originate from within the circuit itself or from DNg100. Within this circuit, E1 is the primary recipient of DNg100 output (fifth-strongest output overall, accounting for 0.49% of DNg100 output synapses). E2 is both the strongest output of E1 and its strongest input (this connection accounts for 2.86% of E1 output synapses and 11.85% of E2 input synapses). I1 is the strongest inhibitory input to E1 (4.57% of E1 input synapses) and the fourth-strongest input to E2 (3.78% of E2 input synapses). This pattern of strong, reciprocal connectivity is what equips the circuit with its rhythm-generating capacity. Importantly, this property would be invisible to conventional connectome analyses: tracing feedforward pathways downstream of DNg100 would identify E1 among the top candidates but miss E2 and I1, while the reverse approach of tracing pathways upstream of motor neurons would miss E1 and I1 because they are not strongly premotor. Path-based graph analyses would fail to recover this core CPG circuit because its function depends on recurrence rather than feedforward connectivity, underscoring the necessity of dynamic simulation for its discovery.

The rhythm-generating capacity of this circuit can be explained quantitatively by performing an eigendecomposition of the dynamical system derived from a linearization of our rate model. The E1-E2-I1 circuit model has one complex-conjugate pair of eigenvalues with an oscillation frequency of ∼14 Hz (**Extended Data Fig. 4c**), which is within the behaviorally relevant range of walking frequencies. Among all 21,544 instances of the E-E-I motif identified in the MANC front leg network, the E1-E2-I1 circuit has the highest intrinsic frequency (**Extended Data Fig. 4f**), further distinguishing it as an outlier in terms of both connection strength and dynamic properties. To confirm that our results do not rely on the specific assumptions of our rate model, we simulated the same circuit using leaky integrate-and-fire neurons and found similar patterns of rhythmic spiking activity (**Extended Data Fig. 5**). This result demonstrates that the rhythm-generating mechanism is a robust property of the circuit’s synaptic architecture rather than an artifact of the modeling framework.

The E1-E2-I1 circuit was sufficient, and both E1 and E2 were necessary, to produce leg motor rhythms in response to DNg100 activation. Motor rhythmicity scores of the three-neuron circuit were comparable to the full VNC network across independent simulation replicates (**Fig. 3i**), and silencing either E1 or E2 in the intact network abolished all rhythmic activity across all four datasets (**Fig. 3j**). This necessity is consistent with the near-universal presence of E1 and E2 across pruning screen replicates, legs, and connectome datasets (**Fig. 3c–f, Extended Data Figs. 6** and **7**). The circuit required strong inhibition, but the identity of the inhibitory cell varied across pruning replicates; neither I1 nor I2 was individually necessary within the full network (**Fig. 3j**). This suggests that the inhibitory neurons I1 and I2 serve partially redundant inhibitory functions within the CPG as part of a larger circuit that coordinates flexible walking rhythms; more candidate cells are listed in **Supplementary Tables 3–6**. One difference between the three-neuron circuit and the full network is that increasing DNg100 input magnitude did not modulate oscillation frequency in the reduced circuit (**Extended Data Fig. 4a**), indicating that additional premotor interneurons in the full network contribute to frequency scaling beyond the core rhythm-generating mechanism.

The minimal three-neuron CPG circuit is repeated in the neuropils of all six legs across all four connectome datasets (**Extended Data Fig. 8**), establishing that this putative rhythm-generating motif is a conserved feature of leg motor circuitry in *Drosophila*. Because no neuron in our model possessed intrinsic bursting or other nonlinear membrane properties, the oscillations arose entirely from synaptic connectivity, consistent with a network oscillator mechanism. The convergence of pruning screens across stochastic replicates, multiple legs, both sexes, and four independent connectome datasets provides strong computational evidence that E1 and E2, together with at least one inhibitory neuron, form the core of a CPG circuit necessary for generating rhythmic leg motor activity during forward walking.

### Motor rhythms in all six legs

Having identified a conserved CPG circuit in each leg neuropil, we next asked whether activating DNg100 bilaterally could drive coordinated rhythmic activity across all six legs. Whether interleg coordination patterns rely from central feedforward mechanisms, sensory feedback, or both remains an open question.

To address this, we scaled up our simulations to the entire MANC connectome (23,532 neurons, including all six leg neuropils, DN axons, sensory axons, and VNC regions associated with the wings, halteres, and abdomen). Bilateral activation of the two DNg100 axons produced oscillatory activity in the core CPG neurons and a subset of motor neurons of all six legs (**Fig. 4a**), demonstrating that a single descending pathway is sufficient to recruit rhythm-generating circuits throughout the nerve cord. Motor rhythms were somewhat less robust in the hind legs and overall rhythmicity was reduced compared to the front leg subnetwork (**Fig. 4b**, *n*=128 replicates), which may reflect differences in connectome reconstruction quality across leg neuropils in the MANC dataset or a difference in the relative strength of feedforward DNg100 drive to these neuropils.

**Figure 4.**
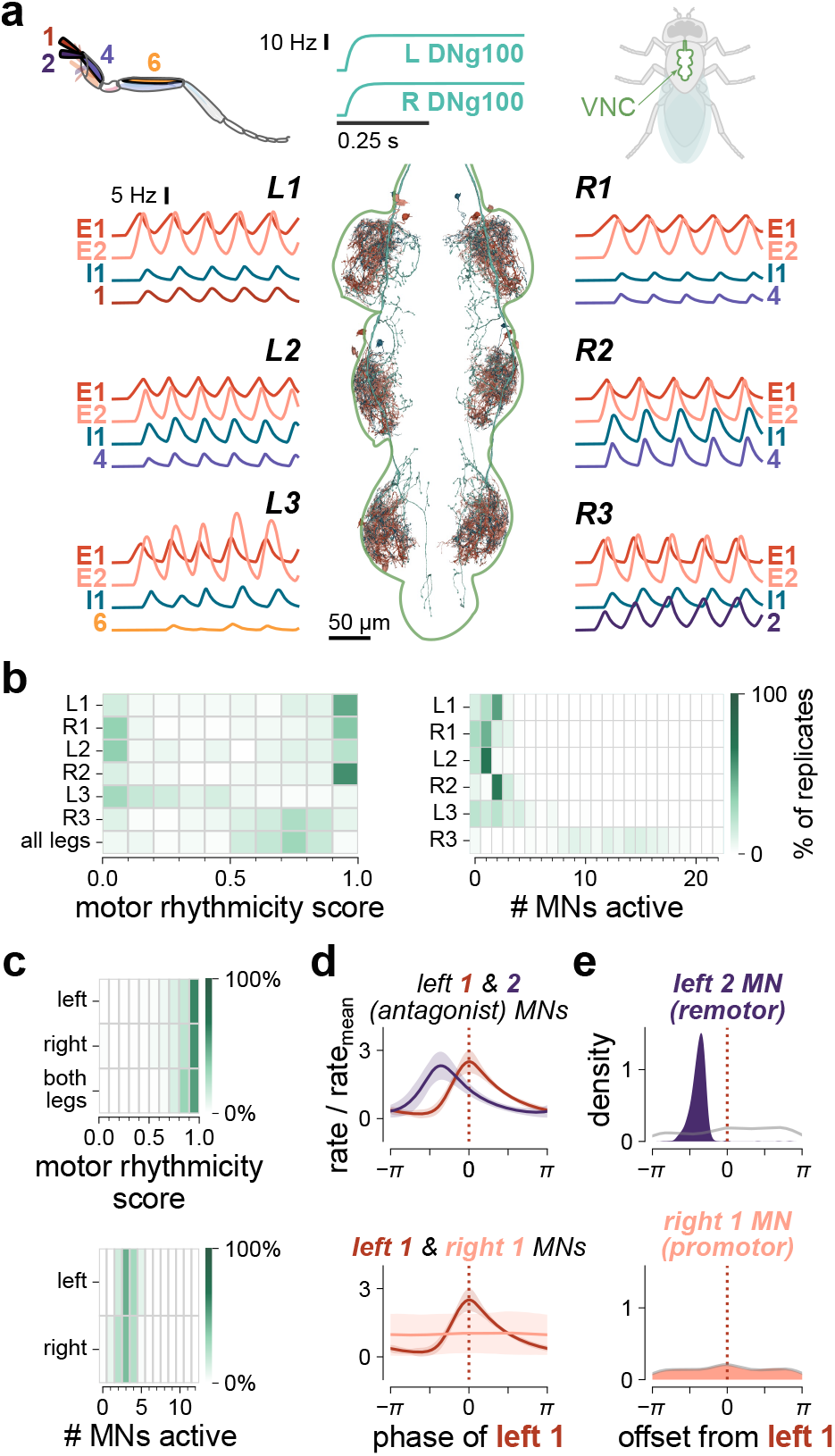
DNg100 drives rhythmic motor activity in all legs in full connectome. **a**, Simulating the full MANC connectome dataset, activating the two DNg100 neurons drove oscillations in the core CPG neurons and a subset of motor neurons in all 6 legs. Anatomy of the core CPG circuit neurons in all 6 leg neuropils (MANC), dorsal view. **b**, Bilateral DNg100 activation drove oscillatory MN activity in all legs, though least consistently in L3 (*n*=128 replicates). The active MN histogram (*right*) excluded 10 replicates where the total number of active motor neurons across legs exceeded 150. **c**, Bilateral DNg100 activation consistently drove oscillatory MN activity in both front legs of the MANC front leg network (*n*=1024 replicates), recruiting a comparable number of motor neurons per leg. **d**, In the front legs network, when both DNg100s were activated, left coxa promomotor rhythmicity score # MNs active tor and remotor MNs displayed a reliable phase offset (*top*), but left vs right coxa promotor motor neurons showed no consistent phase relationship (*bottom*). Plots show mean ± std of MN rates (*n*=1024 replicates) after aligning to the peaks of MN 1 (left coxa promotor). **e**, Distributions (KDE) of phase offsets of the left coxa remotor MN (*top*) and right coxa promotor MN (*bottom*) relative to MN 1 across 1024 replicates, calculated from the peak of the cross-correlation relative to the oscillation period. Gray lines show trial-shuffled control distributions.

Since bilateral DNg100 activation consistently produced robust oscillations in motor neurons of both front legs (**Fig. 4c**), we characterized the phase coordination within and between legs in the MANC front leg subnetwork. Within each leg, we observed a consistent phase offset between antagonistic muscles (coxa promotor and remotor; **Fig. 4d,e**). However, despite robust rhythmic activity in both legs, no consistent phase relationship emerged between the left and right coxa promotor motor neurons (**Fig. 4d,e**), and phase coupling was absent across the six leg CPGs in the full connectome simulation (**Extended Data Fig. 9b**). These results suggest that descending drive from DNg100 alone is sufficient to generate within-leg motor coordination but not realistic interleg coordination. Additional mechanisms, such as phasic sensory feedback or biomechanical coupling, may be required to maintain consistent phase relationships across legs. Identifying the cells and circuits responsible for interleg coordination remains an important open question that our connectome modeling framework is well positioned to address in future work.

### A separate class of descending neurons recruits an overlapping rhythm generating circuit

Having identified the core CPG circuit downstream of DNg100, we returned to examine other high-scoring DNs from our initial computational screen. One cell type, DNb08, stood out because all four of its neurons (two per side) produced among the highest rhythmicity scores in the screen (**Fig. 1e, Supplementary Table 1**). Although a study had created a genetic driver line for DNb08 neurons [54], their function had not been previously investigated. Activating a single DNb08 neuron in the full network simulation consistently produced rhythmic leg motor neuron activity across three of the four connectome datasets (MANC, FANC, and mCNS; **Fig. 5a,b**). To test this prediction experimentally, we optogenetically activated DNb08 neurons in decapitated flies and found that they reliably produced rhythmic leg movements (**Fig. 5c–e, Supplementary Video 2**). However, these movements differed qualitatively from the coordinated walking driven by DNg100 stimulation—they resembled the searching movements that insects exhibit when their legs are not in contact with the ground [55]. Indeed, movements were evoked more reliably when the spherical treadmill was removed. Overall, these results confirmed a model prediction about a previously unstudied descending pathway, and demonstrate that connectome simulations can reveal the motor function of uncharacterized cell types without prior experimental knowledge.

**Figure 5.**
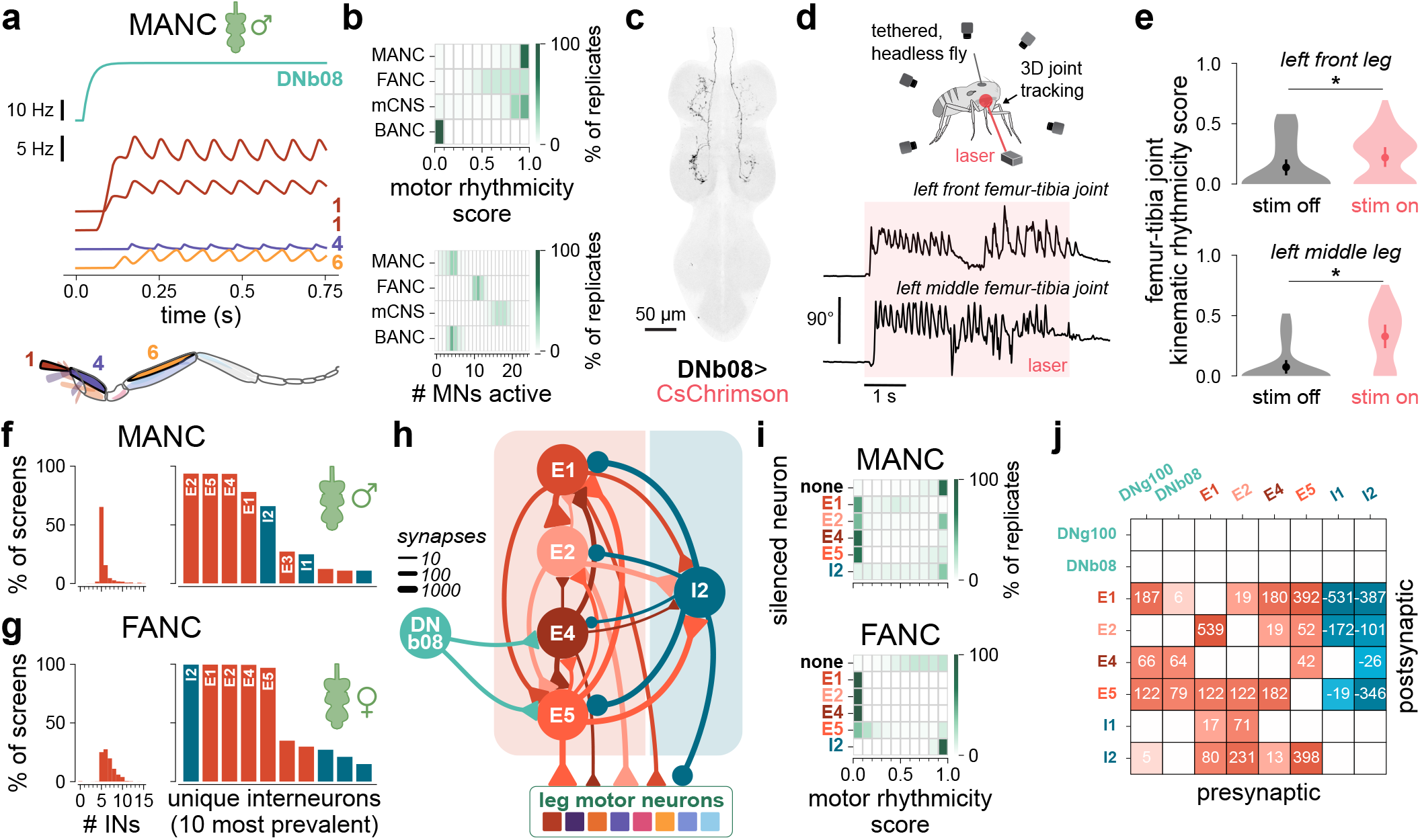
DNb08 produces rhythmic leg movements in simulations and headless flies, and its downstream network converges to an overlapping minimal circuit. **a**, Activation of DNb08 in the full MANC network produced rhythmic leg motor activity. All active motor neurons are shown. Muscle innervation of MNs is indicated on traces. **b**, DNb08 activation consistently produced leg motor rhythms in MANC, FANC, and mCNS, though not the BANC (*n*=1024 replicates each). **c**, VNC expression of Split-GAL4 line labeling DNb08 neurons (SS70620*>*CsChrimson). **d**, Femur-tibia joint kinematics in front and middle legs during optogenetic activation of DNb08 neurons in a headless fly (see **Supplementary Video 2**). **e**, Activating DNb08 produced oscillatory movement of the front and middle legs (*n*=10 flies, 8 stim-on and 4 stim-off trials per fly). Mean and error bars (95% c.i.) computed across means of individual flies (*p* = 0.036, 0.004 for paired *t*-tests comparing stimulus on vs. off trials for front and middle leg, respectively). Violin plots show overall distributions over all trials. **f**, Minimal circuits identified by pruning in MANC. *Left*: Number of interneurons within minimal circuits across 1024 independent simulations (37 screens did not converge to 15 or fewer interneurons and are not shown). *Right*: Percentage of screens in which minimal circuits contained specified interneurons (see **Supplementary Table 7** for full list of cell identities). **g**, Same as **f**, but for FANC (54 screens did not converge to 15 or fewer interneurons and are not shown). See **Supplementary Table 8** for full list of cell identities. **h**, Connectivity of the minimal rhythm-generating circuit downstream of DNb08 (MANC). Triangles and circles indicate excitatory and inhibitory connections, respectively. Synapse counts are summed across all left leg motor neurons. **i**, In both MANC and FANC network simulations, individually silencing E1, E2, E4, and E5 abolished or significantly degraded leg motor neuron rhythmicity, but silencing I2 did not. **j**, Synaptic weight matrix for all cells in either DNg100 and DNb08 minimal circuits (MANC) reveals strong shared circuitry (E1 and E2) and a common motif of excitatory-inhibitory connections.

To isolate the core circuit generating rhythmic activity downstream of DNb08, we repeated our computational sufficiency screen. The most common minimal circuit, identified in nearly half (45.4%) of the 1024 replicates, consisted of five interneurons (**Fig. 5f,h, Supplementary Table 7**): the same E1, E2, and I2 neurons from the core walking CPG, plus two additional excitatory neurons “E4” (IN03A006) and “E5” (INXXX464). Unlike DNg100, which drives E1 directly, DNb08 is only weakly presynaptic to E1. Instead, E4 and E5 receive strong input from DNb08 and relay excitation to E1. I2 then provides strong feedback inhibition onto all four excitatory interneurons, implementing the same delayed inhibition mechanism that underlies rhythm generation in the DNg100 circuit (**Extended Data Fig. 10c**). As with the DNg100 circuit, this subcircuit is strongly self-connected, with 8.90% of its total input synapses originating from within the subcircuit or from DNb08. Replication of the pruning screen in FANC converged to the same E1-E2-E4-E5-I2 circuit as the most common solution (246/1024 replicates; **Fig. 5g, Supplementary Table 8**), and silencing any of the four excitatory neurons abolished rhythmic motor patterns in the otherwise intact network while silencing I2 alone did not (**Fig. 5i**). A second convergent circuit, sharing E2, E4, and E5 with E3 and I1, had the same motif and oscillated by the same mechanism (**Extended Data Fig. 10c–f**). Interestingly, E4 and E5 receive substantially more mechanosensory input than E1 and E2 do, providing a potential explanation for why DNb08-driven movements were more reliably evoked when the flies were removed from the treadmill (**Extended Data Fig. 7c**). Taken together, the convergence between the DNg100 and DNb08 minimal oscillating circuits (**Fig. 5j**) reveals that descending neurons supporting distinct behaviors can recruit overlapping VNC subnetworks that share a rhythm-generating core. Thus, the putative CPG we identify here may serve as a common circuit for multiple rhythmic leg behaviors.

## Discussion

By simulating the dynamics of *Drosophila* VNC connectomes, we identified a putative CPG circuit for walking and other rhythmic leg movements. This CPG is downstream of the walking command neuron DNg100 and its core consists of three neurons, two excitatory and one inhibitory (E1-E2-I1), that form a coherent, recurrent circuit (**Fig. 3g**). Each DNg100 neuron synapses onto more than 1,400 cells in the VNC, and the strong recurrence among these downstream circuits make it prohibitively difficult to pinpoint a rhythm generating circuit using traditional connectivity analyses. We instead isolated the core CPG circuit using a computational sufficiency screen that iteratively pruned the VNC network downstream of DNg100 while preserving leg motor neuron rhythms. This circuit was able to produce walking motor rhythms in each of the six legs and in four independent connectome datasets, spanning both sexes. We also found that a separate descending pathway, DNb08, drives rhythmic leg movements through a five-neuron circuit that shares the same two excitatory neurons (E1 and E2) identified downstream of DNg100.

The interneurons identified by our pruning screens receive input from other walking-related DNs, including DNg97 (oDN1, [50]) and DNg74 (web, [56]), as well as grooming-related neurons DNg12 [57] and DNg62 (aDN1, [58]). A full list of descending inputs to the core CPG neurons is provided in **Supplementary Table 9** and motor rhythmicity scores for known movement-related DNs are reported in **Supplementary Table 10**. Because our DN screen works as a discovery tool, we do not interpret low scores as evidence against any pathway’s role in pattern generation, for reasons described above. To further probe the role of the core CPG in rhythm generation, we repeated the DN activation screen with E1 neurons silenced and found that many high-scoring DNs showed significant decreases in motor rhythmicity (**Extended Data Fig. 1d**). This suggests that E1 is a critical node for rhythmic motor output across multiple descending pathways, not only those driving walking. We therefore hypothesize that several rhythmic limbed behaviors share the same core CPG in each leg. Rather than switching among multiple non-overlapping CPGs, the fly VNC may generate diverse motor patterns by modulating the frequency and phase of a shared core circuit, sometimes recruiting additional neurons to transition among behavior-specific outputs. This idea is consistent with past work showing that pattern-generating circuits are multifunctional [59, 60], in that the same neurons participate in the generation of different behaviors.

Our simulation results make several testable predictions that motivate future experiments. Here, we carried out feasible experiments using existing genetic reagents, which confirmed that DNg100 activity controls walking speed (**Fig. 2i,j**) and that activation of DNb08 drives rhythmic leg movements (**Fig. 5d,e**). Testing the remaining predictions will require the creation of new genetic driver lines that specifically label the core CPG interneurons. Chief among these predictions is that optogenetic silencing of E1 and/or E2 will abolish DNg100-driven leg motor rhythms. In contrast, we predict there exist multiple inhibitory cells (e.g., I1 and I2) that are each sufficient to produce rhythmic inhibitory feedback in the CPG network. Because the cells in our model lacked intrinsic bursting, plateau potentials, and post-inhibitory rebound properties, we predict the core CPG operates as a network oscillator. Direct electrophysiological recordings will be required to confirm or refute these intrinsic properties and corroborate our modeling predictions. Targeted recordings and silencing experiments will also help clarify the differential contributions of I1, I2, and other neurons identified in our screens (**Supplementary Tables 3–8**) to the flexible generation of walking rhythms.

### Limitations and future extensions of the model

Activating DNg100 neurons in our connectome simulations recruited motor neurons that innervate muscles throughout the fly leg, and some antagonistic muscle pairs were activated with naturalistic phase relationships. For example, the coxa promotor and remotor muscles, which swing the leg anteriorly during swing phase and posteriorly during stance, respectively, were active with a phase offset resembling that recorded in other walking insects [5]. However, other muscles likely active during walking, including the main tibia flexors [45], were either silent or non-rhythmic in our simulations. This suggests that although the feedforward signals from the CPG circuit may be sufficient to initiate cyclic stepping movements, the naturalistic pattern of muscle activation likely relies on additional factors: proprioceptive feedback, mechanical coupling, neuromodulation, or combinatorial activity of multiple DNs. This interpretation is consistent with extensive evidence that CPG rhythms and sensory feedback act cooperatively during locomotion [6, 61]. One promising direction for future work is to integrate the DNg100 pathway with inputs to the 13A and 13B neuron classes, which coordinate rhythmic leg movements during grooming and receive proprioceptive feedback [62], potentially revealing how sensory context shapes muscle recruitment patterns.

The differential sensitivity to sensory load may also explain a key behavioral difference between the two descending pathways. DNb08-driven movements were more consistent when fly’s legs were unloaded, whereas DNg100 reliably drove walking on the spherical treadmill. This is consistent with the observation that E4 and E5 receive substantially more direct mechanosensory input than E1 and E2 (**Extended Data Fig. 7c**), suggesting that the DNb08 circuit is more sensitive to load-dependent suppression. Future neuromechanical models incorporating proprioceptive feedback dynamics will be important for testing this hypothesis, given the well-established role of load sensing in insect walking and posture control [63, 64, 65]. The behavioral differences between DNg100 and DNb08 stimulation may also reflect their distinct natural functions: DNg100 is a known walking command neuron, but DNb08 may be involved in leg searching, aggression, or grooming. For instance, DNb08 was recently identified as a descending output of the aggression-associated neuron AVLP491 in the male brain [66]. Although both DNs recruit parts of the same core CPG circuit, their downstream circuits differ in their motor outputs: DNb08 recruits the tibia extensor but not the coxa remotor motor neuron (**Fig. 5a**), consistent with a behavior that engages the leg differently than forward walking.

Our key findings replicate across all four connectome datasets, but we observed some differences (**Fig. 2e, Fig. 4b, Fig. 5b**). Most notably, DNb08 failed to drive rhythmic activity in one of the four datasets. We also saw differences in the identities of recruited motor neurons, both across datasets of the same leg and across leg neuropils within the same dataset. These differences likely reflect multiple factors, including variation in motor neuron reconstruction fidelity and synapse detection (**Extended Data Figs. 4g** and **9a**, also see [67]). These factors are difficult to untangle within any single dataset. However, the availability of several biological replicates suggests a future strategy. Comparing simulated dynamics across datasets could support more principled approaches to parameter estimation, such as tuning the synaptic scaling parameter *b* separately for each dataset or neuropil.

In the full DNg100 network model, the frequency of the motor rhythms of the legs increased with higher magnitudes of DNg100 activation—a prediction that we confirmed experimentally using optogenetic manipulations in behaving flies. This frequency scaling was absent in the minimal three-neuron CPG circuit, suggesting that additional interneurons in the full network contribute to frequency modulation beyond the core rhythm-generating mechanism. One hypothesis is that graded recruitment of additional interneurons shifts the drive onto the core CPG, modulating the effective delays in the recurrent circuit that set the oscillation period. Increasing the rate of DNb08 activity did not produce frequency scaling (**Extended Data Fig. 10a**), indicating that input-frequency modulation is not a general property of all descending pathways that recruit the core CPG. Identifying the interneurons responsible for frequency scaling in the DNg100 pathway is an important open question. Extending our computational sufficiency screen to investigate frequency modulation will be a priority for future work.

Our DNg100 simulations did not produce the tripod interleg coordination pattern characteristic of hexapod walking. Several VNC neurons connect the left and right CPG circuits disynaptically, but their inclusion was insufficient to couple the phase of the left and right legs. This suggests that proprioceptive feedback, biomechanical coupling, or other neural pathways may be necessary to organize interleg coordination. It remains an open question which of these mechanisms, or other modifications to the model, would be sufficient to achieve coordinated walking. One promising path forward is to couple VNC connectome simulations to biomechanical models of the fly body interacting with a simulated physical environment [68, 69], which would enable closed-loop testing of how proprioceptive feedback, biomechanical coupling, and muscle dynamics contribute to interleg coordination. Additionally, DNs appear to be recruited as populations, due to high levels of interconnectivity in the brain [70]. Future models could therefore incorporate DN-to-DN connectivity now available in the full CNS connectome datasets [30, 31].

### Theory of minimal rhythm-generating networks

In a neural network of threshold-linear units, a three-neuron circuit with two excitatory cells and one inhibitory cell is a mathematically minimal model for rhythm generation. This follows from early theoretical work by Amari [71], who showed that a limit cycle solution emerges within a regime of synaptic weights between a pair of cells, one excitatory and one inhibitory. In this minimal two-neuron circuit, the excitatory cell must excite itself (i.e., via autapses). Because neurons in the fly CNS rarely form autapses [28, 29], a minimum of two cells that excite each other (e.g., E1, E2) is required to replace this self-excitation [72]. Inhibitory feedback onto the excitatory cells then completes the oscillatory loop, producing alternating activity across the circuit. Indeed, all minimal circuits identified by our pruning screens followed this E-E-I architecture. Beyond minimal rhythm-generating networks, larger inhibition-dominated circuits can in principle support more flexible motor patterns, including distinct gaits [73] and variable oscillatory network dynamics [74].

The mathematical mechanism by which the three-neuron CPG circuit generates rhythms has a striking parallel in oscillatory networks of coupled biological components beyond the nervous system, including evolved and synthetic gene circuits. Circadian rhythms in cyanobacteria, for example, are maintained by three genes that form a pacemaker through positive and negative feedback interactions [75]. In synthetic biology, the *repressilator* produces oscillations using three transcriptional regulators arranged in a cyclic inhibitory loop [76]. Another two-regulator synthetic gene circuit achieves tunable periodicity through a self-activating unit that recruits delayed inhibitory feedback [77]. In our three-neuron circuit, the E1 →E2 connection is crucial in amplifying DNg100 excitation, and E2→ I1 introduces a further time delay before inhibition. In each case, oscillations arise from a balance of excitatory amplification and time-delayed inhibition. That such similar oscillatory motifs have been found in neural circuits and gene regulatory networks suggests that this architecture may represent a principle of biological rhythm generation.

### Connectome simulations as a discovery tool

Beyond rhythmic leg movements, our simulation approach provides a general strategy for discovering functional microcircuits in other parts of connectomes. A key consideration is ensuring that model outputs are directly interpretable in terms of behavior. Here, leg motor neuron activity provided a natural readout that enabled direct experimental validation through optogenetics in behaving flies. The robustness of our results across a wide range of biophysical parameters and synaptic weight perturbations was perhaps surprising, but may reflect two complementary factors. First, our criteria for success, rhythmic activity of leg motor neurons, was deliberately permissive, making the results insensitive to precise parameter values. Second, the core CPG circuit may be intrinsically robust, hardwired by evolution to generate reliable rhythms across a wide range of environmental conditions and behavioral contexts [78]. Together, our results suggest that connectome-constrained simulations need not resolve every unknown biophysical parameter to yield meaningful predictions about circuit function.

To investigate ethological behaviors where movements are coordinated over longer timescales than a single walking bout, connectome simulations may need to incorporate other circuit and cellular parameters such as receptor and ion channel expression, neuronal morphology, neuromodulation, and plasticity. Machine learning approaches, such as reinforcement learning, may also help fine-tune connectome simulations to achieve more complex motor patterns and behavioral sequences embodied in a biomechanical fly body model (as was done with artificial neural networks in [79] and [69]). However, training artificial neural network components to fit parameters in connectome-body interfaces carries risks for biological interpretability [80]. Overall, our work shows that even simplified simulations, when pursued in close collaboration with biological experiments, can predict circuit dynamics from static connectomes and are a compelling approach to study the neural control of animal behavior.

## Acknowledgments

We thank Eiman Azim, Steve Brunton, Michael Dickinson, Michael Elowitz, David Perkel, Adriane Otopalik, Simon Sponberg, and members of the Tuthill and Brunton Labs for comments on the manuscript. We thank Anne Sustar for VNC images, Tony Azevedo for advice on motor neuron modeling, David Samy for help with coding, Salil Bidaye for sharing fly stocks, confocal images, and unpublished results, Kenji Doya for pointing us to Amari’s 1978 textbook in Japanese and Tomo Ouchi for assistance with translating it. This work was funded through a Swartz Foundation Fellowship in Theoretical Neuroscience to E.T.T.A.; a Shanahan Family Foundation Fellowship at the Interface of Data and Neuroscience to D.T.; National Institutes of Health grants R01NS102333, R01NS128785, the New York Stem Cell Foundation, and a Pew Biomedical Scholar Award to J.C.T.; NIH grant R01NS145438 to J.C.T. and B.W.B.; Air Force Office of Scientific Research award FA9550-19-1-0386, NIH grants U01NS136507, R01NS136988, and the Richard & Joan Komen University Chair to B.W.B. J.C.T is a New York Stem Cell Foundation–Robertson Investigator.

## Author Contributions

S.M.P., J.C.T., and B.W.B. conceived of the study, designed the analyses, and wrote the manuscript. S.M.P. designed and developed models for connectome simulation, with assistance from J.K.L., and E.T.T.A. implemented software simulations in JAX. G.M.C. collected and analyzed behavioral data. D.T. derived the dynamical systems analysis and designed and implemented several extensions of the model.

## Data and Code Availability

All code to run the connectome simulations, results from computational experiments, and scripts to reproduce the analyses are available on GitHub (https://github.com/smpuglie/Pugliese_cpg_2025).

## Methods

### VNC connectome datasets

Our VNC connectome simulations were constructed based on four published datasets, the male (MANC, [28]) and female (FANC, [29]) adult nerve cord connectome datasets and the male (mCNS, [31]) and female (BANC, [30]) adult central nervous system datasets. Unless stated otherwise, simulation results throughout the manuscript are from the front leg motor subnetwork within the MANC dataset. To select this subset of cells to simulate from the entire MANC dataset, we first selected all front leg motor neurons (class “motor neuron”, subclass “fl”). Next we added all front leg premotor neurons by querying for any neuron that made synaptic outputs onto any of those leg motor neurons. Finally, we added all descending neurons (class “descending neuron”) that made any synaptic outputs onto the front leg premotor neurons. After collecting this initial set of neuron IDs, we filtered for neurons that were proofread and had been assigned a neurotransmitter prediction. In this way, we constructed the front leg motor network **W** (as illustrated in **Fig. 1b**), defined as the set of recurrent connections between any nodes in this set. Each element |**w**_*ij*_| was the number of synapses detected from neuron *j* to neuron *i*. To remove very weak connections and a small number of neurons that made only very weak connections to this network, we imposed a floor of 5 synapses. The sign of **w**_*ij*_ was assigned to be positive if the presynaptic neuron *j* was predicted to be cholinergic and negative if neuron *j* was predicted to be GABAergic or glutamatergic (according to the “predictedNt” property). This front leg motor subnetwork in MANC consisted of 4,604 neurons, making up 57% of all cells in the front leg neuropils and 20% of cells in the entire connectome dataset.

We followed a very similar procedure in the Male CNS (mCNS) dataset, given the shared data infrastructure through *neu*Print [81]. The only differences were that we used the new property of “consensusNt” rather than “predictedNt” to assign neurotransmitter identities and valence, and we only queried synapses inside the VNC regions of interest to exclude descending-descending and descending-ascending connectivity within the brain that was present in this dataset but not the MANC dataset. This gave us a dataset of 4,310 neurons for the mCNS simulations.

To construct our network from the BANC dataset, we selected the same subset of cells, except that we included all descending neurons for simplicity (*n*=1,314). As in the mCNS dataset, we excluded synapses within cells in our subnetwork but outside of the VNC (i.e., in the central brain), in this case by using a bounding box on the coordinates of the VNC within the EM volume. To assign neurotransmitters, we used the “neurotransmitter verified” property within the codex annotations CAVE table wherever it was available for a neuron, and for neurons without a verified neurotransmitter, we used “neurotransmitter predicted” instead, the outcome of their classifier. One small difference between the datasets in the BANC and FANC as opposed to the mCNS datasets is that we did not impose a minimum synapse count of 5, so all detected synapses were included in **W**. The BANC subnetwork we used totaled 4,963 neurons. Unless otherwise noted, we simulated activation of the DNg100 neuron innervating the left leg neuropils. We refer to this as the “left DNg100,” reflecting its target neuropil rather than its soma location in the right hemisphere, as we only consider connectivity within the VNC.

In the FANC connectome simulations, we applied an analogous set of criteria to construct **W**, but we restricted the network to the left front leg because this neuropil is the most thoroughly proofread and annotated [29, 32]. We started with front left leg motor neurons (from motor_neuron_table_v7 CAVE annotation table), then added local premotor neurons (i.e., those with neurites constrained to the front left leg neuropil, from left_t1_local_premotor_table_v6), and the single descending neuron DNg100, for a total of 803 neurons. We added the two left DNb08 neurons to the dataset to produce the results in **Figure 5**. Unlike the other datasets, the FANC dataset does not have an automated classifier to predict the neurotransmitter for each neuron. However, the developmental hemilineage of neurons with somas within the VNC can be determined anatomically and is highly predictive of the neurotransmitter released by each neuron [32]. We used the hemilineage annotations from the corresponding CAVE tables [82] to assign a positive or negative sign to each cell’s output. The only exceptions to this approach were made because of a set of discrepancies we noticed between the FANC and MANC datasets in hemilineage assignments. Specifically, 4 neurons (segIDs 72905112552773752, 72975481296715415, 72975481363813393, 72905112552782067) are grouped with glutamatergic hemilineages in FANC so would be considered inhibitory, but the same cells in MANC (bodyIDs 162543, 11751, 13246, 15115) have an indeterminate hemilineage assignment of “TBD.” In MANC, an independent neurotransmitter classifier predicted that all 4 of these neurons, belonging to types INXXX464 (E5), INXXX466 (E2), and INXXX468, are cholinergic (with probability *>*0.81), so they would be excitatory. Taken together, we reasoned that since there is no agreement on hemilineage assignments and high agreement about the neurotransmitter prediction, these four cells are most likely excitatory. Therefore, we assigned their output synapse weights to be positive. Once the mCNS and BANC datasets were available, we confirmed that they also show the predicted neurotransmitters of these cells to be acetylcholine. Encouragingly, all three datasets with an automatic classifier predict that all 24 of the cells belonging to these 3 types across the 6 legs are cholinergic with high probability.

For our simulations in the full MANC dataset in **Figure 4**, we simulated all neurons whose proofreading status was Traced.

### Connectome simulation models

We developed a rate model to simulate the activity **r**(*t*) of every cell in the network as a function of its synaptic connectivity **W** and external inputs **I**(*t*):

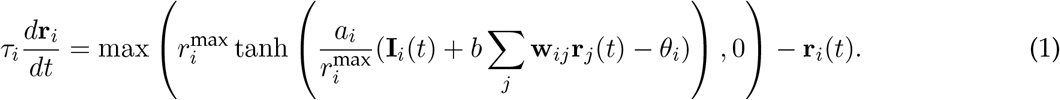

The form of this equation is modified from a standard formulation of a rate model [83]. We report the output of this model as a rate in units of Hz, but as the model itself does not spike, this rate is interpreted as an abstract quantification of activity. The inputs to the DNs in the network are in arbitrary units.

Each cell had four biophysical parameters; below, we elaborate on how these parameters were randomly chosen for each replicate of the simulation. We chose the input nonlinearity to be a positive-rectified hyperbolic tangent function max(tanh(*x*), 0). The function was parameterized as 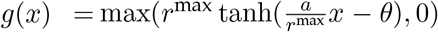 so that parameters have straightforward interpretations. In particular, *r*^max^ is the upper bound, *θ* is the minimal input needed to produce a nonzero output, and *a* is the slope at this threshold value 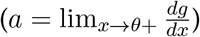(these parameters and their distributions are visualized in **Fig. 1d**). Although this nonlinearity has an upper bound *r*^max^, in most of the rhythm-generating simulation results presented in this paper, the rates stayed well within the approximately linear regime, so that the neural dynamics could have been effectively captured by a simpler nonlinearity (e.g. ReLU). However, we chose the rectified tanh function because some simulations produced oversaturated or unstable output (e.g., in some DN activation screens and noise perturbation shuffles), and capping rates supported more stable numerical solutions with the ODE solver implementation.

In our analyses, we considered neurons to be “active” or “recruited” in a simulation if their rate **r**_*i*_(*t*) exceeds 0.01 Hz at any time point.

#### Biophysical parameter distributions

For each cell and in each random replicate of the simulation, we drew its biophysical parameters from normal distributions truncated at zero, then normalized two of the parameters by neuron size (as elaborated below). Across all simulations, the gain (before size normalization) was *a*^∗^ ∼ 𝒩 (1, 0.1), the threshold (before size normalization) was *θ*^∗^ ∼ 𝒩 (7.5, 0.6), the maximum rate was *r*^max^ ∼ 𝒩 (200, 10) Hz, and the time constant was *τ* ∼ 𝒩 (0.02, 0.002) sec. Next, 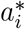 was divided by the median-normalized size of each neuron *i*, and 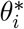 was multiplied by the median-normalized size of each neuron *i*, resulting in distributions for *a* and *θ* with long tails (**Fig. 1d**).

Our choice of these parameter values was based on a combination of physiologically feasible ranges reported in the literature and a hyperparameter search. Since *r*^max^ and *τ* have direct biological interpretations, we used reasonable ranges for rate limits and time constants, respectively, of *Drosophila* neurons given experimental literature [47, 44] and consistency with previous fly neuron simulation studies [48, 37]. In our model, gain was dimensionless and units of input were arbitrary, since voltage was not directly modeled. Therefore, values of *θ* and *a* have no direct biological analogs. In our hyperparameter search, we evaluated a grid of values for the four biophysical parameters as well as the synaptic scaling constant *b* on a set of simple monosynaptic connections. In this test network, we chose ranges of values that produced consistent activity in downstream neurons when stimulated with input and ensured this activity also decayed to quiescence after input was removed.

To normalize *θ* and *a* by neuron size *s*_*i*_, we used morphological data from each dataset. For each cell, *a*^∗^ and *θ*^∗^ were randomly drawn from the distributions described above, then gain was divided by *s*_*i*_ and threshold was multiplied by *s*_*i*_. In MANC and the mCNS, we used the available volume property. In FANC and the BANC, we accessed the neuron meshes in the reconstruction to calculate the surface area of each cell. In all datasets, we took the ratio of each cell’s size to the median cell size in the dataset as the value used to scale gain and threshold. We reasoned that, in most neurons, a significant amount of cell volume is accounted for by the approximately cylindrical neurites, so that volume scales approximately linearly with surface area, which is the determining factor for input resistance. This approach allowed us to account for the general differences in excitability between large and small neurons, and this consideration was critical in our simulation results. Other connectome analyses have normalized the strength of synaptic connections by the total number of synaptic inputs received by the postsynaptic cell [84], which is in practice similar to size normalization, since large neurons tend to receive many synapses. This was a crucial consideration in our model: without adjusting *a* and *θ* for size, the network does not produce robust oscillations in response to DNg100 input even when this input is adjusted down in magnitude to compensate (**Extended Data Fig. 2d**). The descending neurons have larger volumes in the two CNS datasets due to their arbors in the brain being preserved, which is why the DNg100 input needed to be of a higher magnitude in these datasets.

This method of parameter selection was designed to reveal dynamics that arise primarily from synaptic connectivity, without relying on special intrinsic properties or fitted parameters. By drawing parameters independently for each neuron and simulation replicate, we simulated a heterogeneous ensemble of networks sharing the same underlying connectivity structure. To further test the robustness of our rhythmicity scores to the magnitude of the biophysical parameter variance, we repeated our DN activation screen and DNg100 simulations after increasing the standard deviation by 3x for all four of the parameters. Widening the initial distribution of all biophysical parameters decreased the overall motor rhythmicity scores for the top scoring DNs, but the ones we chose to focus on this paper (DNg100 and DNb08) were still among the highest scoring DNs (**Extended Data Fig. 2c**). This widening of variance also led to a larger number of replicates to fail to produce motor rhythms under DNg100 drive (**Extended Data Fig. 2a**), but this drop in mean motor rhythmicity score was modest and did not change the interpretation of our results. About half of the replicates scoring 0 could be attributed to simulations in which there were no motor neurons active. We also performed a sensitivity analysis (**Extended Data Fig. 2b**), restoring one parameter distribution at a time to its previous variance magnitude, and determined that the most sensitive parameter was the gain. In other words, the other 3 parameter distributions could be increased by at least 3x and our DNg100 activation results were nearly unchanged.

We also examined the sensitivity of our results to the synaptic scaling parameter *b* = 0.03 by running simulations across a grid of values, with *b* chosen separately for each of the three neurotransmitter types: *b*_ACh_, *b*_GABA_, and *b*_Glu_ (**Extended Data Fig. 2e**). Increasing *b*_ACh_ to 0.045 still produced viable oscillatory dynamics, but larger deviations resulted in either runaway network activity or insufficient neuron recruitment, both yielding low rhythmicity scores. At *b*_ACh_ = 0.03, a broad range of *b*_GABA_ and *b*_Glu_ values produced stable oscillatory outputs. We expect that jointly adjusting input strength alongside *b*_ACh_ would expand the viable parameter range by compensating for changes in excitatory balance.

#### Implementation of numerical simulations

To compute solutions to the rate models of our connectome simulations, we implemented a numerical differential equation solver using the GPU-accelerated numerical computing library JAX [85]. Our solver used Diffrax [86] with the Runge-Kutta integration method of order 5(4) (diffrax.Dopri5()). Error tolerances were chosen to be rtol=2e-6 and atol=5e-9, based on hyperparameter searches of values that minimized both runtime and sum-squared error when compared to a reference run with very small error tolerances. Run on 4 Nvidia L40s GPUs, 1,024 replicates of the DNg100 activation in the full front leg MANC model typically ran in ∼18 minutes.

#### Leaky integrate-and-fire model

To confirm that the rhythm generating behavior of the core CPG circuit does not depend on our specific rate model formulation, we simulated the dynamics of the same connectome weights using a spiking neuron model. We implemented a leaky integrate-and-fire model based on the same parameter values from previous work in the *Drosophila* brain [37], without tuning or training specific to our circuit or validation against the physiology of these cells. Specifically, we used the following parameters: *V*_rest_ = −52 mV, *V*_reset_ = −52 mV, *V*_thresh_ = −45 mV, *t*_refractory_ = 2.2 ms, *τ*membrane = 20 ms, *τ*_synaptic_ = 5 ms, *w*_synaptic_ = 0.275 mV, *t*_delay_ = 1.8 ms, *V*_init_ = −52 mV, *dt* = 0.01 ms, simulation time = 3 s, network input = 0.15 nA. The membrane time constant was not specifically mentioned in [37], so we computed it based on the membrane resistance *R*_*m*_ = 10 kΩ cm^2^ and membrane capacitance *C*_*m*_ = 2 *µ*F cm^−2^ values reported to arrive at *τ*_membrane_ = *R*_*m*_*C*.

#### Linearized dynamical system analysis

We performed an eigendecomposition of the linearized dynamical system corresponding to the full MANC network and the minimal circuit (core CPG circuit and DNg100) to confirm the oscillatory behavior and oscillation frequencies observed in simulations. For this analysis, we simplified the dynamical system equation to 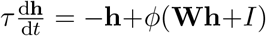, then discretized and linearized it to **h**_*t*+d*t*_ = (1 −*α*+*αg***W**)**h**_*t*_+*αgI*, where *α* = d*t/τ*, and *g* is the non-linearity gain at time *t*, which is potentially different for each neuron (column-wise multiplication with **W**, element-wise multiplication with *I*). With **W**^∗^ ≡ 1 −*α* + *αg***W**, we decomposed the dynamics matrix by eigendecomposition and denoted vectors expressed in the eigen-basis by a hat 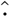. In this basis, we find 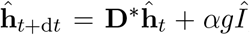, where **D**^∗^ is the diagonal form of **W**^∗^ with eigenvalues *λ*^∗^ = 1 −*α* + *αgλ*.

The system is oscillatory if it has pairs of complex-conjugate eigenvalues. We simplified the solution to find the oscillation frequency of the system for each pair of complex-conjugate eigenvalues. We let 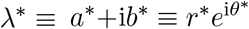, where *θ*^∗^ = atan2(*b*^∗^, *a*^∗^). Thus, we found 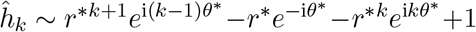, where *k* discretizes time such that *t* = *k*d*t*. We computed the oscillation frequency according to *f* = 1*/*(Δ*k*d*t*), where we chose Δ*k* such that 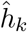 and 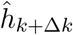 have the same phase. With *α*≪ 1, then *r*^∗^ ≈ 1, and we found Δ*k* ≈ 2*π/θ*^∗^, and finally 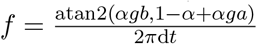 .

To analyze the network dynamics matrix **W** and the core CPG network, we scaled each neuron’s gain according to its size (as described above in ***Biophysical parameter distributions***), then multiplied by an overall gain factor of 0.75 to approximate the simulated nonlinearity (scaled tanh function). Calculations used an average time constant of *τ* = 20 ms and a timestep d*t* = 0.01 ms. Because much of the activity we are interested in is well below saturation (*r*^max^) and in the approximately linear regime of **Eq. 1**, we have not included input **I**(*t*) as a variable in this linearization. For the minimal CPG network, the eigenvalues of **W**^∗^ were {0.9908 + 0.0879*j*, 0.9908 − 0.0879*j*, 0.8683, 0.95}, confirming the oscillatory nature of the system with an oscillation frequency of approximately 14 Hz.

#### Occurrence of oscillator circuit motifs in connectome dataset

We searched for circuit motifs in the VNC connectome that match the structure of the core CPG, namely an inhibitory neuron “I” that inhibits two other excitatory neurons “E_A_” and “E_B_,” among which there is an excitatory chain from “E_A_” to “E_B_” to “I” (**Extended Data Fig. 4e**). We avoided double counting circuits by selecting the circuit with larger sum of synapses in the excitatory chain (“E_A_” → “E_B_” + “E_B_” → “I”). To compare these results with what would be expected of a randomly shuffled network, we permuted the VNC connectome by shuffling all the outgoing connections of each individual neuron, thus preserving the E/I identity of each neuron and its out-degree.

We performed eigenvalue decomposition on these motifs and found that most of them (19,799 out of 21,544, or 91.90%) contain imaginary eigenvalues, in other words are capable of producing oscillations, though most were much slower than the E1-E2-I1 circuit (**Extended Data Fig. 4f**). That most of these motifs are oscillatory is not surprising since the circuit structure is highly likely to produce imaginary eigenvalues. In a shuffle control, across 100 random seeds, there was an average of 302.66 oscillatory motifs out of 306.95 total motifs.

### Motor rhythmicity score

To quantify the degree to which circuit simulations generated rhythmic activity patterns in leg motor neurons, we developed a *motor rhythmicity score* that was used throughout the paper to evaluate and screen circuit models. Our goal was to devise a simple metric, based on autocorrelations, with the property that repeating signals would receive high scores even if they were not necessarily symmetric waveforms (e.g., both a sine wave and a sawtooth would receive a high score).

The motor rhythmicity score assigned to each simulation was computed as the average of rhythmicity scores for all active motor neurons. A motor neuron was included if its **r**_*i*_(*t*) *>* 0.01 at any time after *t* = 250 ms. To compute a motor neuron’s rhythmicity score, we take the trace of its activity after the initial transient response until the end of the simulation (250 ms to end). This trace was normalized so that it is between −1 and 1, 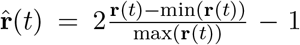. We then computed the autocorrelation of 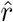 and detected peaks in this autocorrelation, with a prominence threshold of 0.05. The “raw score” for this neuron was assigned as the smaller of the maximum peak height (magnitude above zero) and the maximum peak prominence (magnitude between peak and valley). Using the time shift Δ*t* value at which the most prominent peak occurred as the period of the dominant rhythm, we then computed the same score for a reference sinusoidal waveform with the same frequency and total duration. To normalize the rhythmicity score between 0 and 1, we divided the raw score by the reference score of this frequency-matched sinusoidal signal. Examples of individual motor neuron traces with their scores are shown in **Fig. 1e**. If a motor neuron is active, but there is no prominent peak in the autocorrelation, the neuron received a score of 0.

#### Oscillation frequency and phase analysis

In several analyses, we quantified either the oscillation frequency of simulated neurons (**Fig. 2g, Extended Data Fig. 4a, Extended Data Fig. 10**) or the phase offsets between pairs of oscillating neurons (**Fig. 4e, Extended Data Fig. 9b**). An individual neuron’s oscillation frequency was computed identically to the rhythmicity score calculation: the period *T*_*i*_ = 1*/f*_*i*_ was defined as the time shift (Δ*t*) of the highest peak in the autocorrelation of the normalized activity trace after 250 ms. The overall motor neuron frequency for a simulation was defined as the mean frequency across all oscillating motor neurons. To compute the phase offset of one neuron’s activity **r**_*j*_ relative to another **r**_*i*_, we first calculated the oscillation period *T*_*i*_ of **r**_*i*_ as described above. We then identified the peak of the cross-correlation between **r**_*i*_ and **r**_*j*_ within a time window restricted to 0 ±*T*_*i*_. This time shift Δ*t* at which this peak occurred was normalized to *T*_*i*_ and wrapped to convert it to a phase shift Δ*ϕ*_*j,i*_ ∈ [−*π, π*]. As a control, we assessed the distribution of phase offsets Δ*ϕ*_*j,i*_ across *n* independent replicates by randomly permuting the replicate indices from which the **r**_*j*_ traces were drawn.

#### Kinematic rhythmicity score

To quantify the degree to which leg joint kinematics were rhythmic in DNb08 activation experiments **Fig. 5e**, we used the same calculation as the motor rhythmicity score to evaluate individual joint angle traces. We used the joint angle during a 2 second time window, beginning 0.5 seconds after stimulus onset. Traces for which the detected peak frequency was slower than 1 Hz were assigned a score of 0.

### Descending neuron activation screen

We conducted a computational screen in which each of the 933 excitatory descending neurons (DNs) in our model was stimulated with a tonic input, and we assessed how rhythmic motor neuron outputs driven by that DN were across 128 replicates (as quantified by the motor rhythmicity score). Inhibitory neurons were excluded because they cannot activate other neurons in an otherwise-quiescent network, guaranteeing a score of zero. Each simulation replicate was 1 second in duration, with the tonic input stimulus onset at 20 ms. Because the input strength *I*_stim_ was arbitrary in magnitude in our model, we automatically tuned *I*_stim_ over a range of values depending on if the simulation was underactive or over-saturated. Underactive simulations were defined as having fewer than 5 neurons recruited in the whole network, and oversaturated simulations were defined as having more than 1500 neurons recruited, because such simulations were generally unstable. (In the screen where DNs were coactivated by type, we increased our lower bound on active neurons to 10 neurons.) If the simulation was underactive, *I*_stim_ was doubled, or set to the midpoint to the next highest value that had been tried. If it was oversaturated, *I*_stim_ was halved, or set to the midpoint to the next lowest value that had been tried. In each replicate, this automated adjustment was repeated up to a maximum of 10 times to try to achieve a reasonable stimulus strength. If after these 10 iterations the simulation was still underactive or oversaturated, it was considered uninterpretable and the motor rhythmicity score for this replicate was not computed (set to NaN). Out of the 933 neurons screened, 859 (92.1%) had at least one feasible replicate out of 128, and 845 (90.6%) had at least 50 feasible replicates. Reported motor rhythmicity scores for each neuron (**Fig. 1e, Supplementary Tables 1** and **10**) were the mean score of the feasible replicates.

There were a significantly larger fraction of neurons that had at least one of the 128 replicates score *>*0.5 (233 DNs, representing 27.1% of the DNs with any usable replicates and 25.0% of the DNs overall, **Extended Data Fig. 1b**). Such cases are difficult to interpret and likely include false negatives: DNs that drive biological rhythms but were missed by our screen due to parameter discrepancies with the true biological system, or because our approach specifically identified network oscillators and could not detect rhythms that rely on intrinsic cellular properties such as bursting. We see this not as a weakness of our approach, but rather a feature of using our screen as a circuit discovery tool, much like forward genetic screens are used to identify candidate genes for further investigation. We therefore focused on the small set of neurons that robustly generated motor rhythms across most parameter configurations, rather than over-interpreting the low scores of other neurons. We also show the median and interquartile range to give a sense of the spread of the score distributions among different replicates (**Extended Data Fig. 1a**).

#### Combinatorial descending neuron activation screen

In order to test the network response to differential activation of multiple DNs, we developed an approach to quantify the overall effects of driving randomly sampled combinations of DNs. This combinatorial activation screen explored an orders-of-magnitudes larger space of potential DN coactivations than the one-at-a-time DN screens, so our approach was through stochastic sampling, rather than exhaustive search. We varied the stimulation intensity of each neuron stochastically to sample the large, combinatorial space of activating all 1,318 DNs. Previously, only excitatory DNs were screened because the network had no spontaneous activity; here, inhibitory DNs were included because they can coactivate the network along with excitatory DNs. For each total stimulus amplitude, we split this input stochastically among the DNs. We used a Dirichlet distribution parameterized by a constant parameter *α* to sample combinations of DNs being stimulated. Small values of *α* corresponded to very few DNs stimulated at a time, and larger values of *α* means that many DNs received some input. The single-DN screens effectively corresponded to the limit of *α* →0.

We varied the total stimulus amplitude to cover a few orders of magnitude, from the network barely being active to the network saturating due to the bounded transfer function, and we varied *α* to cover group sizes from a single DNs receiving the majority of the stimulus (*α* = 10^−4^) to almost all DNs sharing a fair share of the total stimulus amplitude (*α* = 10). By running approximately 60,000–80,000 simulations for each point in a grid of total amplitude and *α* parameters, we were able to see that the median motor rhythmicity score for all values in this grid is low, but for moderate levels of input to small numbers of DNs, there are some choices that give high scores (**Extended Data Fig. 1e**). When we examined which neurons contributed the most input overall to the highest-scoring simulations throughout the grid (score *>*0.75), 14/29 of these top rhythmic DNs were also among the highest-scoring DNs in the individual screen, and 2/29 were GABAergic neurons (**Supplementary Tables 1** and **2**).

### Computational sufficiency screen with iterative pruning

We developed an iterative procedure to isolate a subset of cells in the full network that were sufficient to generate rhythmic motor patterns in a reduced model (illustrated in **Fig. 3b**). We started with a full intact network and one random seed for the biophysical parameters of its cells (these parameters were frozen for all iterations in one pruning procedure). As in the DN activation screen, simulations were 1 second in duration and consisted of a tonic input to a single descending neuron with an onset at 20 ms. For DNg100 activation, *I*_stim_ = 250 in MANC, 150 in FANC, and 400 in the two CNS datasets (these activations have arbitrary units and were set to the same values that produced robust oscillations in **Figure 2**). For DNb08 activation, *I*_stim_ = 65 in MANC and 180 in FANC. Leg motor neurons were not considered by the pruning screen, so they all remain present in all simulated networks.

Each replicate of the computational sufficiency screen executed the following stochastic procedure to prune neurons from the model network, until convergence to a minimal network according to the stopping criterion. First, all neurons that were not active were pruned from the network. Thereafter, at each step, one additional neuron was chosen to be silenced, with a probability inversely proportional to its maximum rate at the last iteration. We then simulated DN activation of this new network and evaluated its motor rhythmicity score. If the motor rhythmicity score of this pruned network still exceeded a threshold value (≥0.5), this silenced neuron, along with any inactive neurons, were permanently pruned in the next iteration. If the motor rhythmicity score of a pruned network fell below the threshold, then it was returned to the network. This procedure was then repeated by choosing another neuron to silence. Finally, the stopping criterion was when no neuron in the pruned network could be removed without producing a motor rhythmicity score below threshold, at which point we considered the pruning screen to have converged. The remaining cells in the network formed a subcircuit (with the DN input, recurrent network of interneurons, and all motor neuron outputs) that was sufficient to generate rhythmic activity, in the sense that they were computationally isolated from the initial full network simulation. Because the biophysical parameters and the choices of neurons to silence were stochastic, we repeated the screen 1024 times for each full network activated by each DN (DNg100 and DNb08).

## Fly behavior and optogenetics experiments

### Genotypes used

| Name | Genotype | Figures |
| --- | --- | --- |
| DNg100>CsChrimson | w[1118]/w[1118]; VT058557-GAL4.AD/+;<br>R85F12-GAL4.DBD/20xUAS-CsChrimson-tdTomato<br>su(Hw)attP1 | 2h,i,j<br>ED3 |
| SS70620>CsChrimson | w[1118]/w[1118]; P{y[+t7.7]<br>w[+mC]=R94D12-p65.AD}attP40/+; P{y[+t7.7]<br>w[+mC]=VT031392-GAL4.DBD}attP2/20xUAS-<br>CsChrimson-tdTomato su(Hw)attP1 | 5c,d,e |

#### Optogenetics experiments with 3D joint kinematics

Methods for behavioral experiments were previously described in [44]. Briefly, female flies were cold-anesthetized, de-winged, decapitated, and tethered (0.1 mm tungsten rod) with UV glue (KOA 300). For DNg100 experiments, we used flies that were 10–12 days old, as in [50], because this produced more robust walking behavior, possibly due to accumulated CsChrimson expression. For DNb08 experiments, we used flies that were 2–5 days old. Each fly recovered for 10–15 minutes before being positioned in the behavioral arena. For DNg100 experiments, flies were positioned on an air-supported spherical treadmill (0.13 g, 9.08 mm diameter). To activate DNs, a LED laser (638 nm, 1200 Hz, 30% duty cycle, Laserland) was focused at the body-coxa joint of the front left leg. Each trial consisted of 5 seconds pre-stim, 5 seconds stim-on, and 5 seconds post-stim, for a total trial length of 15 seconds. Trials were run in blocks of 12: 8 experimental trials with the LED laser stimulus on and 4 control trials where the stimulus was off. We recorded each trial using 6 high-speed cameras (300 fps; Basler acA800–510 µm; Balser AG) and tracked 3D leg joint kinematics using DeepLabCut [87] and Anipose [88]. The movements of the ball were recorded with a FLIR camera (FMVU-03MTM-CS, 30 FPS) and processed with FicTrac [89] to read out the forward velocity of the fly. We used a custom Python script to analyze kinematics based on the Anipose tracking data. Stepping frequency was computed on the tracked tarsus tip position along the anterior-posterior axis of the fly during the stimulus period. The frequency of each trial was calculated as the inverse of the average time between peaks, which was detected with scipy.signal.find peaks using a minimum prominence of 0.15 and a minimal temporal distance of 50 ms.

To test DNg100 modulation of walking speed in headless flies, we used laser intensities of 0.03, 0.09, and 0.33 mW/mm^2^ (8 experimental trials of each intensity per fly). DNg100 activation produced forward walking on the ball, but at a lower speed than is typical of wild-type flies [53]. They generally exhibited a tetrapod coordination pattern (**Extended Data Fig. 3**).

## Supplementary Information

**Supplementary Video 1**: Optogenetic activation of DNg100 neurons drives walking in headless flies. Corresponds to experimental data in **Fig. 2h–j**.

**Supplementary Video 2**: Optogenetic activation of DNb08 neurons drives oscillatory leg movements in headless flies. Corresponds to experimental data in **Fig. 5d,e**.

**Supplementary Table 1.** Top scoring neurons in descending neuron activation screen. All descending neurons with a mean motor rhythmicity score *>*0.5, corresponding to **Fig. 1e**.

| bodyId | type | mean motor rhythmicity score |
| --- | --- | --- |
| 18153 | DNp41 | 0.921685 |
| 16683 | DNxl091 | 0.854303 |
| 10093 | DNg100 | 0.850086 |
| 10339 | DNg100 | 0.836744 |
| 30919 | DNg12 | 0.793973 |
| 18279 | DNp41 | 0.791299 |
| 44321 | DNit011 | 0.777426 |
| 32815 | DNg12 | 0.739332 |
| 23461 | DNfl031 | 0.730864 |
| 14061 | DNb08 | 0.727479 |
| 19821 | DNg54 | 0.697184 |
| 37139 | DNit013 | 0.676787 |
| 14966 | DNb08 | 0.657400 |
| 10291 | DNxl134 | 0.651129 |
| 12221 | DNxl121 | 0.643625 |
| 11164 | DNa13 | 0.641784 |
| 25931 | DNa07 | 0.638171 |
| 30130 | DNxn167 | 0.620578 |
| 13892 | DNb08 | 0.618388 |
| 10896 | DNxl133 | 0.615813 |
| 16264 | DNxn159 | 0.604836 |
| 10101 | DNg15 | 0.585706 |
| 23962 | DNxn127 | 0.584200 |
| 11145 | DNa13 | 0.582096 |
| 14680 | DNb08 | 0.575963 |
| 32742 | DNg12 | 0.558370 |
| 22194 | DNxn171 | 0.520236 |
| 10967 | DNa13 | 0.518630 |
| 11930 | DNp43 | 0.516324 |

**Supplementary Table 2.** Highest-contributing neurons to oscillatory simulations in the combinatorial descending neuron screen. All descending neurons which corresponded to ≥0.5% of the input in simulations scoring ≥0.75 in the combinatorial descending neuron screen.

| bodyId | type | predicted neurotransmitter |
| --- | --- | --- |
| 10093 | DNg100 | acetylcholine |
| 10562 | DNxl014 | acetylcholine |
| 14522 | DNp63 | acetylcholine |
| 14680 | DNb08 | acetylcholine |
| 14966 | DNb08 | acetylcholine |
| 16075 | DNp63 | acetylcholine |
| 16541 | DNxl092 | acetylcholine |
| 16683 | DNxl091 | acetylcholine |
| 18153 | DNp41 | acetylcholine |
| 18279 | DNp41 | acetylcholine |
| 19821 | DNg54 | acetylcholine |
| 21158 | DNxn119 | acetylcholine |
| 23461 | DNfl031 | acetylcholine |
| 25645 | DNut054 | acetylcholine |
| 25931 | DNa07 | acetylcholine |
| 30919 | DNg12 | acetylcholine |
| 31408 | DNfl020 | acetylcholine |
| 31531 | DNfl018 | acetylcholine |
| 32742 | DNg12 | acetylcholine |
| 32815 | DNg12 | acetylcholine |
| 33287 | DNnt013 | acetylcholine |
| 33545 | DNfl018 | acetylcholine |
| 34862 | DNg12 | acetylcholine |
| 37139 | DNit013 | acetylcholine |
| 38587 | DNg12 | acetylcholine |
| 44321 | DNit011 | acetylcholine |
| 46922 | DNxn109 | acetylcholine |
| 26318 | DNfl027 | gaba |
| 39216 | DNfl016 | gaba |

**Supplementary Table 3.** Converged minimal circuits for DNg100 pruning screen in MANC. All distinct sets of interneurons that appeared at least 10 times out of 1024 pruning screens.

| bodyIds | types | count |
| --- | --- | --- |
| 10707, 11751, 13905 | IN17A001 (E1), INXXX466 (E2), IN16B036 (I1) | 636 |
| 10707, 10715, 11751, 12026, 17664 | IN17A001 (E1), IN19B012 (E3), INXXX466 (E2), IN13A010 (-), MNfl10 (-) | 123 |
| 10242, 10707, 11751 | IN19A007 (I2), IN17A001 (E1), INXXX466 (E2) | 102 |
| 10707, 10715, 11751, 12026 | IN17A001 (E1), IN19B012 (E3), INXXX466 (E2), IN13A010 (-) | 33 |
| 10707, 11751, 11799, 17664, 162543 | IN17A001 (E1), INXXX466 (E2), IN19A020 (-), MNfl10 (-), INXXX464 (E5) | 11 |

**Supplementary Table 4.** Converged minimal circuits for DNg100 pruning screen in FANC. All distinct sets of interneurons that appeared at least 10 times out of 1024 pruning screens.

| segIds | types | count |
| --- | --- | --- |
| 72483380721505294, 72834674820722635, 72975481296715415, 73820799177797612 | IN19A007 (I2), IN17A001 (E1), INXXX466 (E2), IN19B012 (E3) | 721 |
| 72483380721505294, 72834674820722635, 72905112552773752, 72975481296715415 | IN19A007 (I2), IN17A001 (E1), INXXX464 (E5), INXXX466 (E2) | 89 |
| 72694075576809644, 72834674820722635, 72975481296715415, 73820799177797612 | IN13A010 (-), IN17A001 (E1), INXXX466 (E2), IN19B012 (E3) | 36 |
| 72483380721505294, 72834674820722635, 72905112552773752, 72975481296715415, 72975481364113281 | IN19A007 (I2), IN17A001 (E1), INXXX464 (E5), INXXX466 (E2), 8A (-) | 25 |
| 72834674820722635, 72905112619913957, 72975481296715415, 73820799177797612 | IN17A001 (E1), IN16B036 (I1), INXXX466 (E2), IN19B012 (E3) | 15 |
| 72483380721505294, 72834674820722635, 72975481296715415, 73891167921933971 | IN19A007 (I2), IN17A001 (E1), INXXX466 (E2), 19B (+) | 13 |
| 72483380721505294, 72834674820722635, 72905112552773752, 72975481296715415, 73468887542677672 | IN19A007 (I2), IN17A001 (E1), INXXX464 (E5), INXXX466 (E2), 12B (-) | 13 |

**Supplementary Table 5.** Converged minimal circuits for DNg100 pruning screen in mCNS. All distinct sets of interneurons that appeared at least 10 times out of 1024 pruning screens.

| bodyIds | types | count |
| --- | --- | --- |
| 800173, 800374, 800863 | IN17A001 (E1), IN19A007 (I2), INXXX466 (E2) | 655 |
| 800173, 800863, 801884 | IN17A001 (E1), INXXX466 (E2), IN16B036 (I1) | 181 |
| 800173, 800216, 800863, 809385, 902491 | IN17A001 (E1), IN19B012 (E3), INXXX466 (E2), IN13A010 (-), IN19A020 (-) | 61 |
| 800173, 800216, 800863, 809385 | IN17A001 (E1), IN19B012 (E3), INXXX466 (E2), IN13A010 (-) | 22 |
| 800173, 800216, 800863, 902491 | IN17A001 (E1), IN19B012 (E3), INXXX466 (E2), IN19A020 (-) | 17 |
| 800173, 800216, 800863, 801568, 902491 | IN17A001 (E1), IN19B012 (E3), INXXX466 (E2), IN19A024 (-), IN19A020 (-) | 13 |
| 800173, 800216, 800863, 801568, 809385 | IN17A001 (E1), IN19B012 (E3), INXXX466 (E2), IN19A024 (-), IN13A010 (-) | 10 |

**Supplementary Table 6.** Converged minimal circuits for DNg100 pruning screen in the BANC. All distinct sets of interneurons that appeared at least 10 times out of 1024 pruning screens.

| segIds | types | count |
| --- | --- | --- |
| 720575941469024064, 720575941504247575, 720575941569601650 | INXXX466 (E2), IN17A001 (E1), IN19A007 (I2) | 681 |
| 720575941469024064, 720575941504247575, 720575941544954556 | INXXX466 (E2), IN17A001 (E1), IN16B036 (I1) | 231 |
| 720575941469024064, 720575941504247575, 720575941542442076, 720575941569601650 | INXXX466 (E2), IN17A001 (E1), IN19B012 (E3), IN19A007 (I2) | 36 |
| 720575941469024064, 720575941504247575, 720575941520072883, 720575941542442076 | INXXX466 (E2), IN17A001 (E1), IN19A020 (-), IN19B012 (E3) | 18 |
| 720575941469024064, 720575941504247575, 720575941569601650, 720575941571519483 | INXXX466 (E2), IN17A001 (E1), IN19A007 (I2), INXXX464 (E5) | 15 |

**Supplementary Table 7.** Converged minimal circuits for DNb08 pruning screen in MANC. All distinct sets of interneurons that appeared at least 10 times out of 1024 pruning screens.

| **bodyIds** | **types** | **count** |
| --- | --- | --- |
| 10242, 10707, 11751, 12021, 162543 | IN19A007 (I2), IN17A001 (E1), INXXX466 (E2), IN03A006 (E4), INXXX464 (E5) | 465 |
| 10715, 11751, 12021, 13905, 162543 | IN19B012 (E3), INXXX466 (E2), IN03A006 (E4), IN16B036 (I1), INXXX464 (E5) | 129 |
| 10707, 11751, 12021, 13905, 17664, 162543 | IN17A001 (E1), INXXX466 (E2), IN03A006 (E4), IN16B036 (I1), MNfl10 (-), INXXX464 (E5) | 26 |
| 10707, 11751, 12021, 13905, 162539, 162543 | IN17A001 (E1), INXXX466 (E2), IN03A006 (E4), IN16B036 (I1), IN13A001 (-), INXXX464 (E5) | 21 |
| 10707, 11751, 12021, 13905, 162543 | IN17A001 (E1), INXXX466 (E2), IN03A006 (E4), IN16B036 (I1), INXXX464 (E5) | 12 |
| 10242, 10707, 11751, 21162, 162543 | IN19A007 (I2), IN17A001 (E1), INXXX466 (E2), IN20A.22A036 (+), INXXX464 (E5) | 12 |
| 10242, 10707, 11751, 12021, 21162, 162543 | IN19A007 (I2), IN17A001 (E1), INXXX466 (E2), IN03A006 (E4), IN20A.22A036 (+), INXXX464 (E5) | 10 |

**Supplementary Table 8.** Converged minimal circuits for DNb08 pruning screen in FANC. All distinct sets of interneurons that appeared at least 10 times out of 1024 pruning screens.

| segIds | types | count |
| --- | --- | --- |
| 72483380721505294, 72624462142676168, 72834674820722635, 72905112552773752, 72975481296715415 | IN19A007 (I2), IN03A006 (E4), IN17A001 (E1), INXXX464 (E5), INXXX466 (E2) | 246 |
| 72483380721505294, 72624462142673647, 72624462142676168, 72834674820722635, 72905112552773752, 72975481296715415 | IN19A007 (I2), 3A (+), IN03A006 (E4), IN17A001 (E1), INXXX464 (E5), INXXX466 (E2) | 84 |
| 72413150020127131, 72483380721505294, 72624462142676168, 72834674820722635, 72905112552773752, 72975481296715415 | 22A (+), IN19A007 (I2), IN03A006 (E4), IN17A001 (E1), INXXX464 (E5), INXXX466 (E2) | 76 |
| 72483380721505294, 72624462142676168, 72834674820722635, 72905112552773752, 72905181339304184, 72975481296715415 | IN19A007 (I2), IN03A006 (E4), IN17A001 (E1), INXXX464 (E5), 8A (-), INXXX466 (E2) | 46 |
| 72413150020127131, 72483380721505294, 72624462142676168, 72834674820722635, 72905112552773752, 72905181339304184, 72975481296715415 | 22A (+), IN19A007 (I2), IN03A006 (E4), IN17A001 (E1), INXXX464 (E5), 8A (-), INXXX466 (E2) | 25 |
| 72483380721505294, 72624462142676168, 72834674820722635, 72905112552773752, 72905112619846398, 72975481296715415 | IN19A007 (I2), IN03A006 (E4), IN17A001 (E1), INXXX464 (E5), 8A (-), INXXX466 (E2) | 25 |
| 72413150020127131, 72483380721505294, 72624462142673647, 72624462142676168, 72834674820722635, 72905112552773752, 72975481296715415 | 22A (+), IN19A007 (I2), 3A (+), IN03A006 (E4), IN17A001 (E1), INXXX464 (E5), INXXX466 (E2) | 24 |
| 72483380721505294, 72624462142673647, 72624462142676168, 72834674820722635, 72905112552773752, 72905181339304184, 72975481296715415 | IN19A007 (I2), 3A (+), IN03A006 (E4), IN17A001 (E1), INXXX464 (E5), 8A (-), INXXX466 (E2) | 22 |
| 72413150020127131, 72483380721505294, 72624462142676168, 72834674820722635, 72905112552773752, 72905112619846398, 72975481296715415 | 22A (+), IN19A007 (I2), IN03A006 (E4), IN17A001 (E1), INXXX464 (E5), 8A (-), INXXX466 (E2) | 12 |
| 72483380721505294, 72624462142676168, 72834674820722635, 72905112552773752, 72905112619846398, 72905181339304184, 72975481296715415 | IN19A007 (I2), IN03A006 (E4), IN17A001 (E1), INXXX464 (E5), 8A (-), 8A (-), INXXX466 (E2) | 11 |
| 72483380721505294, 72624462142673647, 72624462142676168, 72834674820722635, 72905112552773752, 72905112619846398, 72975481296715415 | IN19A007 (I2), 3A (+), IN03A006 (E4), IN17A001 (E1), INXXX464 (E5), 8A (+), INXXX466 (E2) | 11 |
| 72483380721505294, 72624462142673647, 72624462142676168, 72624462142690635, 72834674820722635, 72905112552773752, 72975481296715415 | IN19A007 (I2), 3A (+), IN03A006 (E4), 3A (+), IN17A001 (E1), INXXX464 (E5), INXXX466 (E2) | 10 |
| 72483380721505294, 72624462142676168, 72624462142690635, 72834674820722635, 72905112552773752, 72975481296715415 | IN19A007 (I2), IN03A006 (E4), 3A (+), IN17A001 (E1), INXXX464 (E5), INXXX466 (E2) | 10 |
| 72483380721505294, 72624462142673647, 72624462142676168, 72834674820722635, 72905112552773752, 72975481296715415, 72976099705187525 | IN19A007 (I2), 3A (+), IN03A006 (E4), IN17A001 (E1), INXXX464 (E5), INXXX466 (E2), 12B (-) | 10 |

**Supplementary Table 9.** Top descending inputs to DNg100 core CPG circuit interneurons. Table of all descending neurons that make ≥10 synaptic connections onto left front leg E1, E2, and I1 neurons.

| pre.<br>bodyId | post.<br>bodyId | pre.<br>type | pre.<br>nickname | post.<br>type | post.<br>nickname | neuro-<br>transmitter | num. of<br>synapses |
| --- | --- | --- | --- | --- | --- | --- | --- |
| 10093 | 10707 | DNg100 | BDN2 | IN17A001 | E1 | acetylcholine | 187 |
| 10097 | 11751 | DNxl080 |  | INXXX466 | E2 | gaba | 173 |
| 10532 | 10707 | DNg74 | web | IN17A001 | E1 | gaba | 160 |
| 10086 | 11751 | DNg105 |  | INXXX466 | E2 | gaba | 126 |
| 10941 | 11751 | DNxl130 |  | INXXX466 | E2 | acetylcholine | 93 |
| 10279 | 10707 | DNxl049 |  | IN17A001 | E1 | acetylcholine | 92 |
| 23461 | 10707 | DNfl031 |  | IN17A001 | E1 | acetylcholine | 86 |
| 27936 | 13905 | DNfl023 |  | IN16B036 | I1 | acetylcholine | 82 |
| 10107 | 11751 | DNg74 | web | INXXX466 | E2 | gaba | 80 |
| 10291 | 10707 | DNxl134 |  | IN17A001 | E1 | acetylcholine | 75 |
| 19287 | 10707 | DNfl036 |  | IN17A001 | E1 | acetylcholine | 73 |
| 27523 | 10707 | DNfl025 |  | IN17A001 | E1 | acetylcholine | 73 |
| 10420 | 10707 | DNg93 |  | IN17A001 | E1 | gaba | 72 |
| 22875 | 13905 | DNg12 |  | IN16B036 | I1 | acetylcholine | 68 |
| 10058 | 11751 | DNg108 |  | INXXX466 | E2 | gaba | 62 |
| 15249 | 10707 | DNg17 |  | IN17A001 | E1 | acetylcholine | 60 |
| 156265 | 10707 | DNfl014 |  | IN17A001 | E1 | acetylcholine | 54 |
| 10291 | 11751 | DNxl134 |  | INXXX466 | E2 | acetylcholine | 40 |
| 21235 | 10707 | DNfl032 |  | IN17A001 | E1 | gaba | 39 |
| 32742 | 10707 | DNg12 |  | IN17A001 | E1 | acetylcholine | 38 |
| 32815 | 10707 | DNg12 |  | IN17A001 | E1 | acetylcholine | 37 |
| 11220 | 10707 | DNxl127 |  | IN17A001 | E1 | gaba | 36 |
| 20782 | 10707 | DNfl034 |  | IN17A001 | E1 | acetylcholine | 34 |
| 24753 | 10707 | DNg62 | aDN1 | IN17A001 | E1 | acetylcholine | 34 |
| 31078 | 10707 | DNg12 |  | IN17A001 | E1 | acetylcholine | 33 |
| 10656 | 10707 | DNg97 | oDN1 | IN17A001 | E1 | acetylcholine | 32 |
| 21320 | 13905 | DNfl033 |  | IN16B036 | I1 | acetylcholine | 22 |
| 14903 | 10707 | DNg17 |  | IN17A001 | E1 | acetylcholine | 21 |
| 10107 | 13905 | DNg74 |  | IN16B036 | I1 | gaba | 20 |
| 10107 | 10707 | DNg74 |  | IN17A001 | E1 | gaba | 19 |
| 27350 | 10707 | DNxl070 |  | IN17A001 | E1 | gaba | 19 |
| 14836 | 13905 | DNxl099 |  | IN16B036 | I1 | acetylcholine | 19 |
| 27516 | 10707 | DNxn143 |  | IN17A001 | E1 | acetylcholine | 18 |
| 10058 | 10707 | DNg108 |  | IN17A001 | E1 | gaba | 17 |
| 10532 | 11751 | DNg74 |  | INXXX466 | E2 | gaba | 16 |
| 31635 | 10707 | DNg12 |  | IN17A001 | E1 | acetylcholine | 16 |
| 13845 | 13905 | DNg44 |  | IN16B036 | I1 | glutamate | 15 |
| 28478 | 10707 | DNg12 |  | IN17A001 | E1 | acetylcholine | 15 |
| 13257 | 10707 | DNxl103 |  | IN17A001 | E1 | acetylcholine | 15 |
| 19287 | 13905 | DNfl036 |  | IN16B036 | I1 | acetylcholine | 15 |
| 23953 | 10707 | DNfl030 |  | IN17A001 | E1 | acetylcholine | 14 |
| 11354 | 10707 | DNg75 |  | IN17A001 | E1 | acetylcholine | 13 |
| 13257 | 13905 | DNxl103 |  | IN16B036 | I1 | acetylcholine | 12 |
| 25879 | 10707 | DNfl029 |  | IN17A001 | E1 | acetylcholine | 12 |
| 27217 | 10707 | DNfl024 |  | IN17A001 | E1 | acetylcholine | 12 |
| 10760 | 10707 | DNa01 |  | IN17A001 | E1 | acetylcholine | 12 |
| 10410 | 11751 | DNd03 |  | INXXX466 | E2 | glutamate | 12 |
| 11270 | 11751 | DNxl129 |  | INXXX466 | E2 | acetylcholine | 11 |
| 21320 | 10707 | DNfl033 |  | IN17A001 | E1 | acetylcholine | 11 |
| 17547 | 10707 | DNfl040 |  | IN17A001 | E1 | acetylcholine | 11 |
| 17400 | 10707 | DNfl040 |  | IN17A001 | E1 | acetylcholine | 11 |
| 10410 | 10707 | DNd03 |  | IN17A001 | E1 | glutamate | 11 |
| 11961 | 13905 | DNxl126 |  | IN16B036 | I1 | gaba | 10 |
| 16369 | 10707 | DNfl015 |  | IN17A001 | E1 | acetylcholine | 10 |

**Supplementary Table 10.** Mean motor rhythmicity scores of selected known motor-related descending neurons in activation screen. DNg97 is the standard cell type of oDN1 [50]. DNg62 is the standard cell type of aDN1 [58].

| bodyId | type | mean motor rhythmicity score |
| --- | --- | --- |
| 10093 | DNg100 | 0.850086 |
| 10339 | DNg100 | 0.836744 |
| 30919 | DNg12 | 0.793973 |
| 32815 | DNg12 | 0.739332 |
| 14061 | DNb08 | 0.727479 |
| 14966 | DNb08 | 0.657400 |
| 13892 | DNb08 | 0.618388 |
| 14680 | DNb08 | 0.575963 |
| 32742 | DNg12 | 0.558370 |
| 38587 | DNg12 | 0.481810 |
| 35473 | DNg12 | 0.426328 |
| 13809 | MDN | 0.401343 |
| 14419 | MDN | 0.391900 |
| 18107 | DNa11 | 0.284322 |
| 11070 | DNg97 | 0.252115 |
| 35413 | DNg12 | 0.176919 |
| 31613 | DNg12 | 0.144966 |
| 29120 | DNg12 | 0.090160 |
| 33586 | DNg12 | 0.089108 |
| 30346 | DNg12 | 0.083595 |
| 30999 | DNg12 | 0.076487 |
| 22870 | DNg12 | 0.065393 |
| 34862 | DNg12 | 0.061884 |
| 34382 | DNg12 | 0.061433 |
| 31123 | DNg12 | 0.060159 |
| 10656 | DNg97 | 0.056609 |
| 40338 | DNg12 | 0.051124 |
| 31078 | DNg12 | 0.042315 |
| 31742 | DNg12 | 0.039892 |
| 31635 | DNg12 | 0.038536 |
| 10126 | DNa02 | 0.037617 |
| 18953 | DNg08 | 0.037082 |
| 23518 | DNg08 | 0.035260 |
| 26725 | DNg12 | 0.033749 |
| 31361 | DNg12 | 0.030724 |
| 32424 | DNg12 | 0.029012 |
| 24753 | DNg62 | 0.026326 |
| 28478 | DNg12 | 0.015591 |
| 27728 | DNg12 | 0.011012 |
| 25606 | DNg08 | 0.009654 |
| 152849 | DNg62 | 0.008469 |
| 32667 | DNg12 | 0.006763 |
| 10118 | DNa02 | 0.003438 |
| 17155 | DNa11 | 0.002761 |
| 14523 | MDN | 0.001584 |
| 34253 | DNg12 | 0.001121 |
| 29706 | DNg12 | 0.000658 |
| 12955 | DNp09 | 0.000427 |
| 12926 | DNp09 | 0.000000 |
| 30271 | DNg12 | 0.000000 |
| 13438 | MDN | 0.000000 |
| 24458 | DNg08 | 0.000000 |
| 22875 | DNg12 | 0.000000 |
| 24858 | DNg12 | 0.000000 |

**Extended Data Fig 1.**
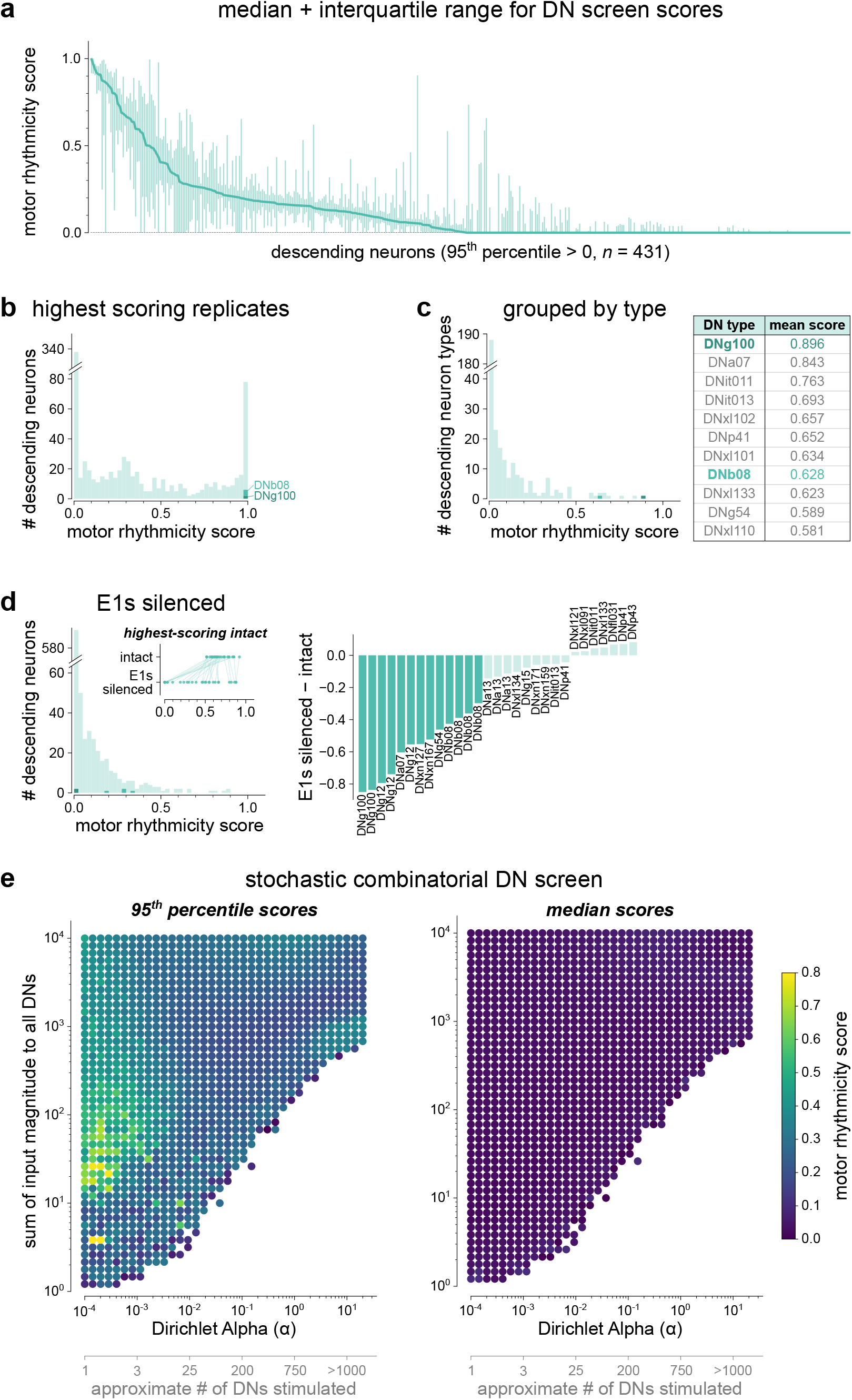
Additional description of descending neuron screen results. **a**, Median and interquartile range for the DNs which had a 95th percentile score above 0 (*n*=128 replicates). **b**, Histogram of the best score out of the 128 replicates for each excitatory DN. **c**, Results of a DN screen procedure where all cells of the same type were coactivated with an equal input (*n*=16 replicates, 321 excitatory DN types). Table shows the types that had an average score ≥0.5. As in other analyses, scores do not distinguish whether active motor neurons belong to one or both legs. **d**, Results of a DN screen procedure where both E1 (IN17A001) neurons were silenced (*n*=16 replicates). The inset and bar plot show the change in scores for the top scoring neurons in the intact screen (**Supplementary Table 1**). **e**, Results of the stochastic combinatorial screen (see Methods). Simulations with no motor neuron activity are removed. Secondary *x*-axis gives a heuristic interpretation of the Dirichlet *α* parameter.

**Extended Data Fig 2.**
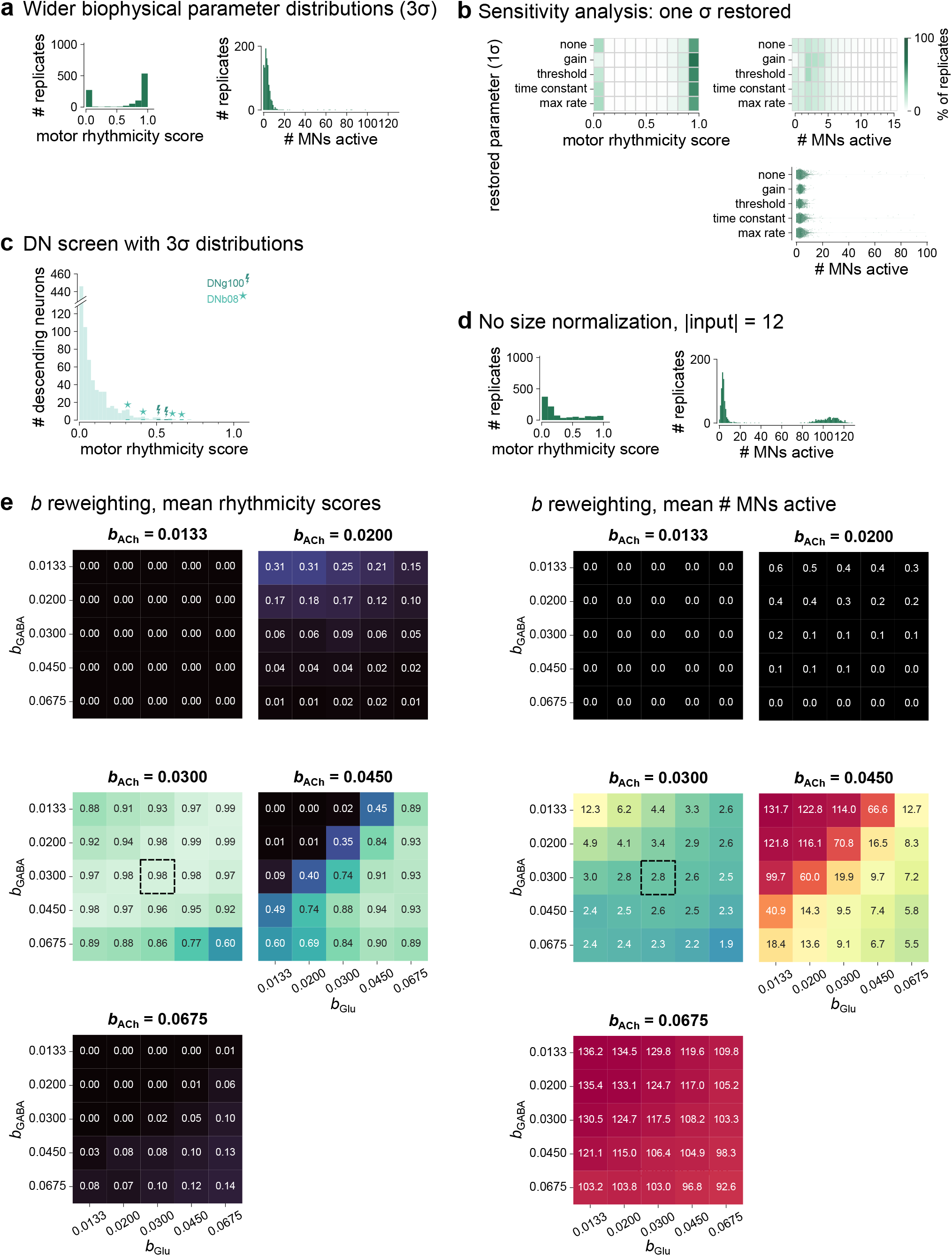
Further exploration of model hyperparameters. **a**, Results of the DNg100 activation simulations in networks with a 3-fold increase in standard deviations of all four parameters (gain, threshold, time constant, and maximum rate), *n*=1024 replicates. **b**, DNg100 with 3-fold increases in standard deviations of all but one parameter, *n*=1024 replicates each with same random seed. The results are most sensitive to gain, which restores consistent performance when it is returned to its original distribution. Rescuing time constant and maximum rate have no effect. **c**, Results of the descending neuron screen in networks with a 3-fold increase in standard deviations of all four parameters DN screen, *n*=16 replicates. **d**, Results of the DNg100 activation simulations in which the size normalization was removed and the input to DNg100 was reduced to an amplitude of 12. **e**, Exploration of the effects of varying separate weighting parameters *b* for each of the three neurotransmitters in the model to model potential systematic differences in their synaptic efficacy.

**Extended Data Fig 3.**
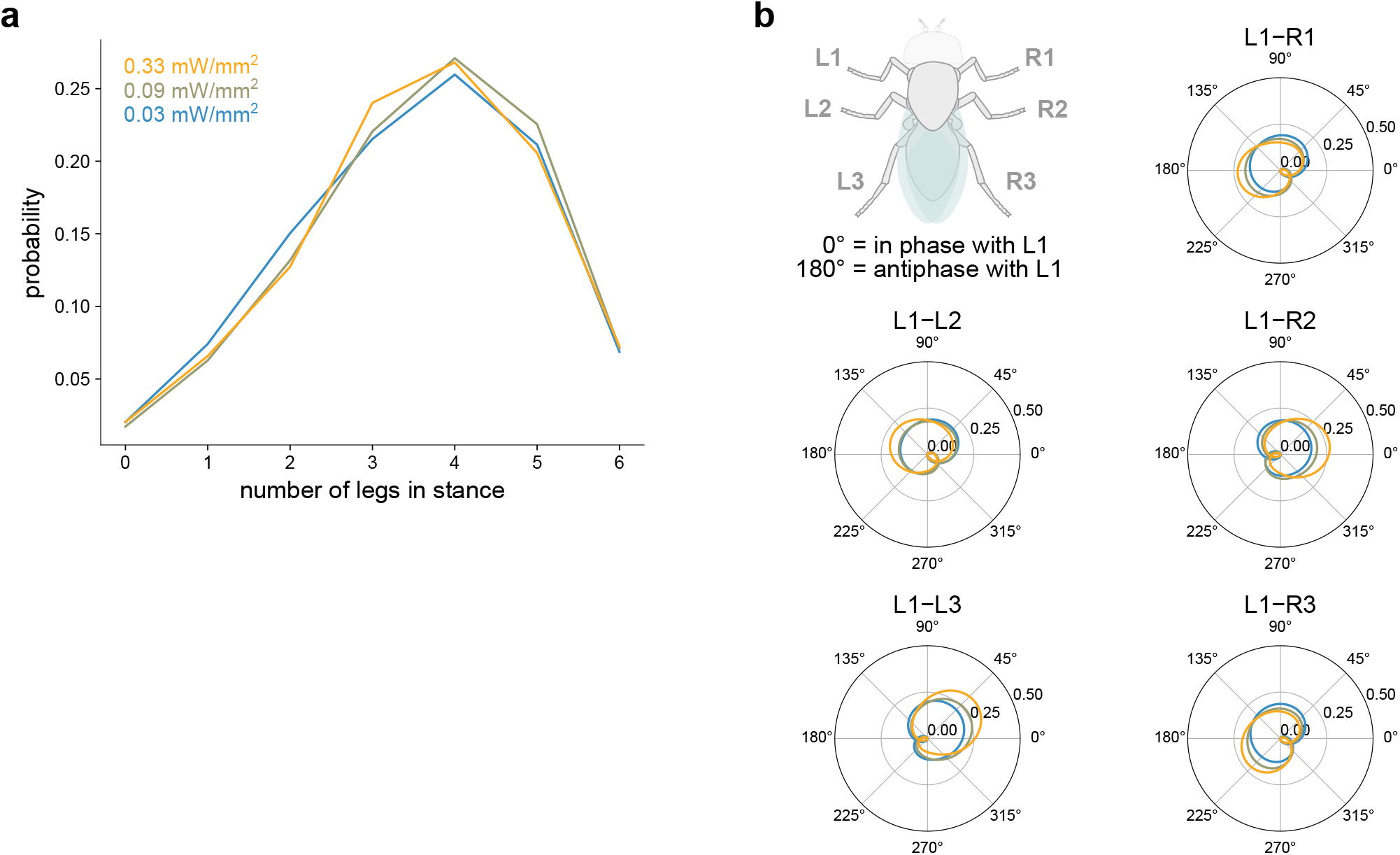
Interleg coordination patterns during optogenetic DNg100 activation in headless flies. **a**, Probability of *n* legs being in stance simultaneously during DNg100-driven walking, with *n* from 0 to all 6 legs, at each laser intensity. **b**, Relative phase between legs at each laser intensity.

**Extended Data Fig 4.**
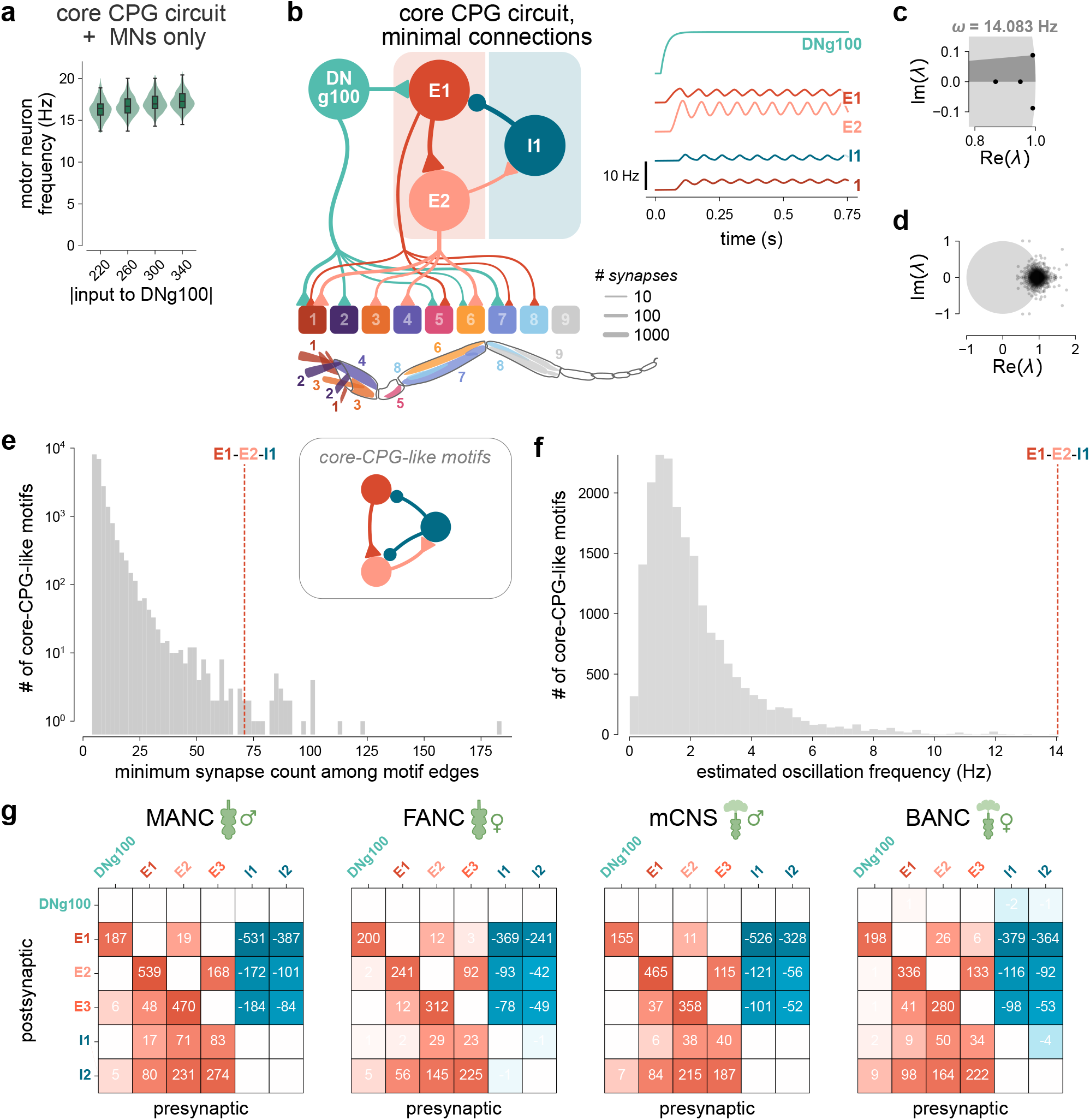
Additional details of the core CPG circuits. **a**, The MANC core CPG circuit does not show the same modulation of motor neuron frequency with DNg100 input strength as the full network does (*n*=512 replicates per condition). Simulations with no active motor neurons were omitted. Green shaded regions are distributions of frequencies, and black bars indicate medians and quartiles. **b**, MANC core CPG circuit in which all interneuron-interneuron connections aside from the cyclic E1 →E2 →I1 →E1 connections have been removed still produces rhythmic activity. **c**, Eigenvalues of the linearized dynamical system of the 4-neuron subcircuit consisting of only the left DNg100, and the front left E1, E2, and I1. The light gray region denotes the unit circle in the complex plane. Darker gray region highlights the angle to the complex eigenvalue pair, which corresponds to the phase advance per time step. This pair of complex-valued eigenvalues corresponds to an oscillation frequency of about 14 Hz. **d**, Eigenvalues of the linearized dynamical system of the full MANC front leg motor network. **e**, Histogram of the weakest of the four required connections out of the core-CPG-like 3-neuron motifs in the MANC front leg subnetwork. **f**, Histogram of the fundamental frequencies of each of the oscillatory core-CPG-like 3-neuron motifs in the MANC front leg subnetwork. **g**, Connectivity of the E1, E2, E3, I1, and I2 neurons in the front left leg neuropil of all four connectome datasets.

**Extended Data Fig 5.**
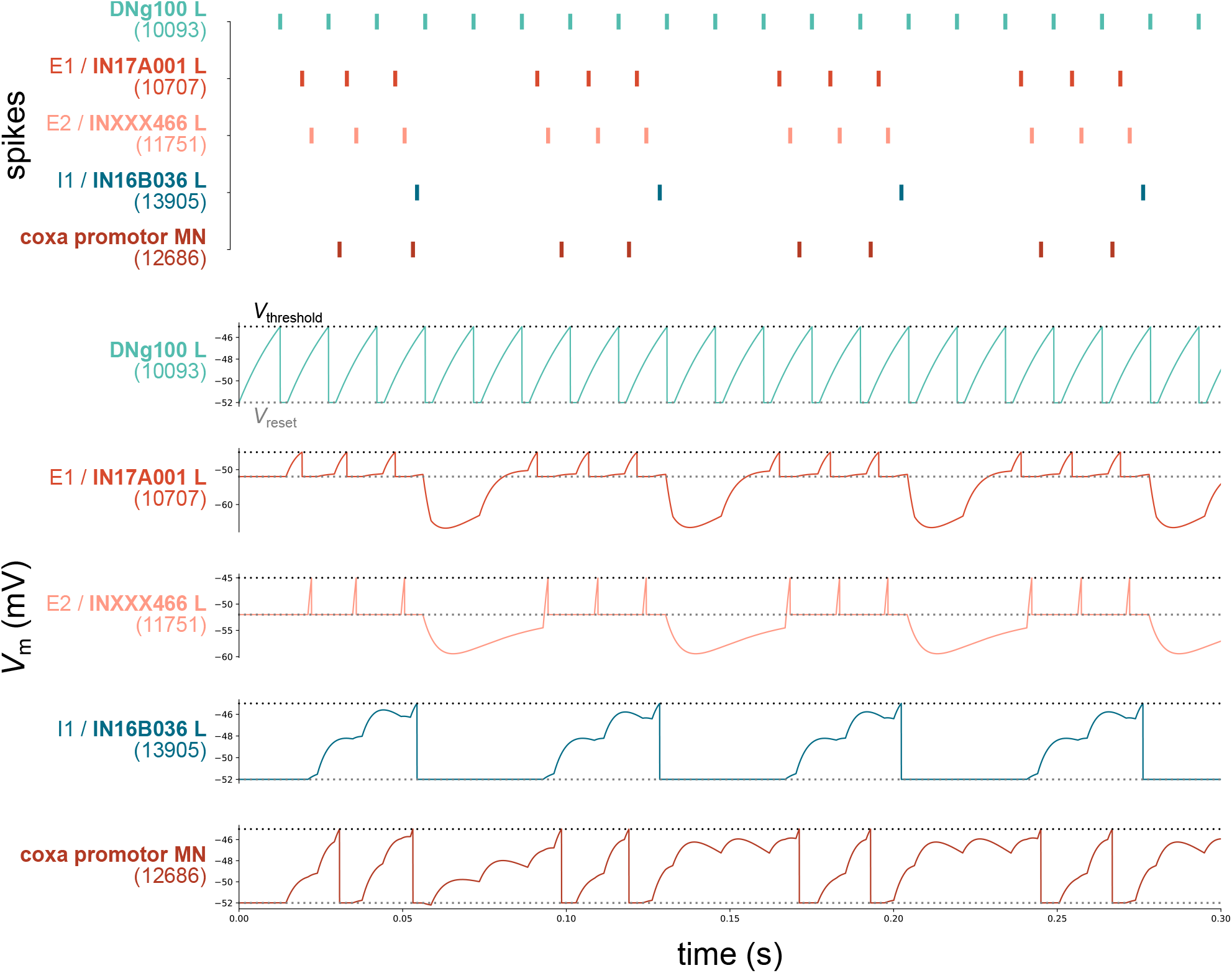
Leaky integrate-and-fire model of core CPG circuit showed rhythmic activity. Simulation included a subcircuit consisting of only the left DNg100, E1 L, E2 L, I1 L, and all of leg motor neurons. *Top*: Spike times, and *Bottom*: voltage (*V*_m_) for each of the core CPG neurons, and one motor neuron.

**Extended Data Fig 6.**
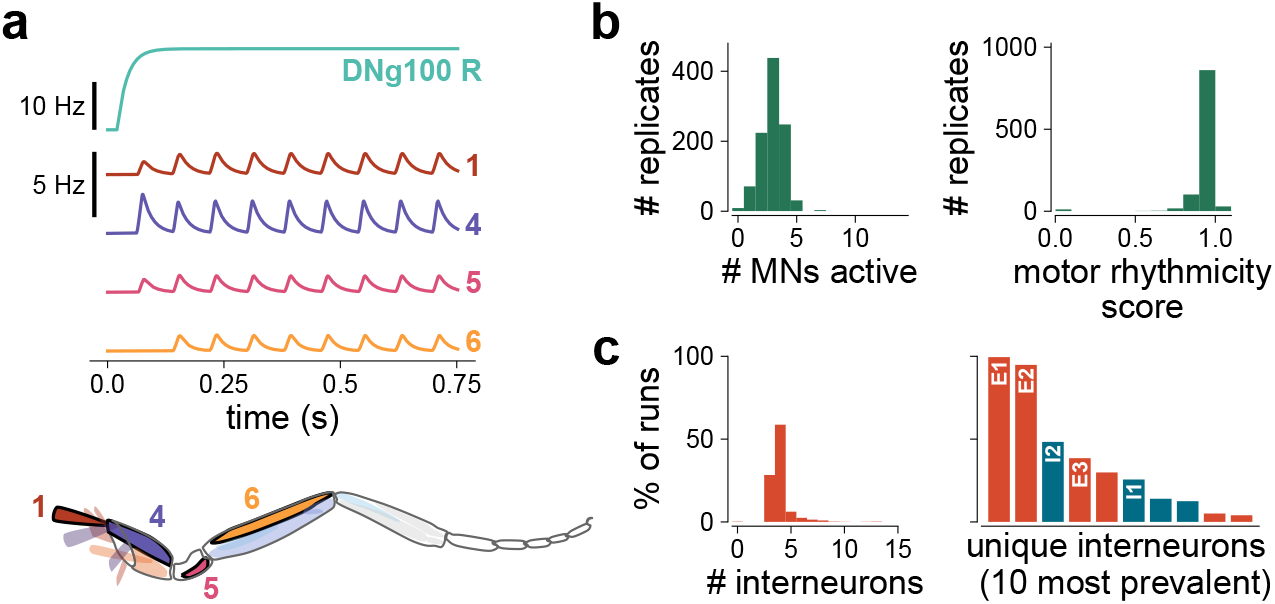
DNg100 R activation. **a**, Example activity of MANC network with step DNg100 R stimulation. Selected motor neurons shown, with numbers corresponding to muscles innervated in the anatomical diagram. **b**, DNg100 R activation consistently produces motor rhythms (*n*=1024 replicates). **c**, Circuits identified by pruning (*n*=1024 replicates, right DNg100, MANC). *Left*: Number of interneurons that are in minimal circuits (14 screens did not converge to 15 or fewer interneurons and are not shown). *Right*: Percentage of simulation runs in which minimal circuits contained unique interneurons.

**Extended Data Fig 7.**
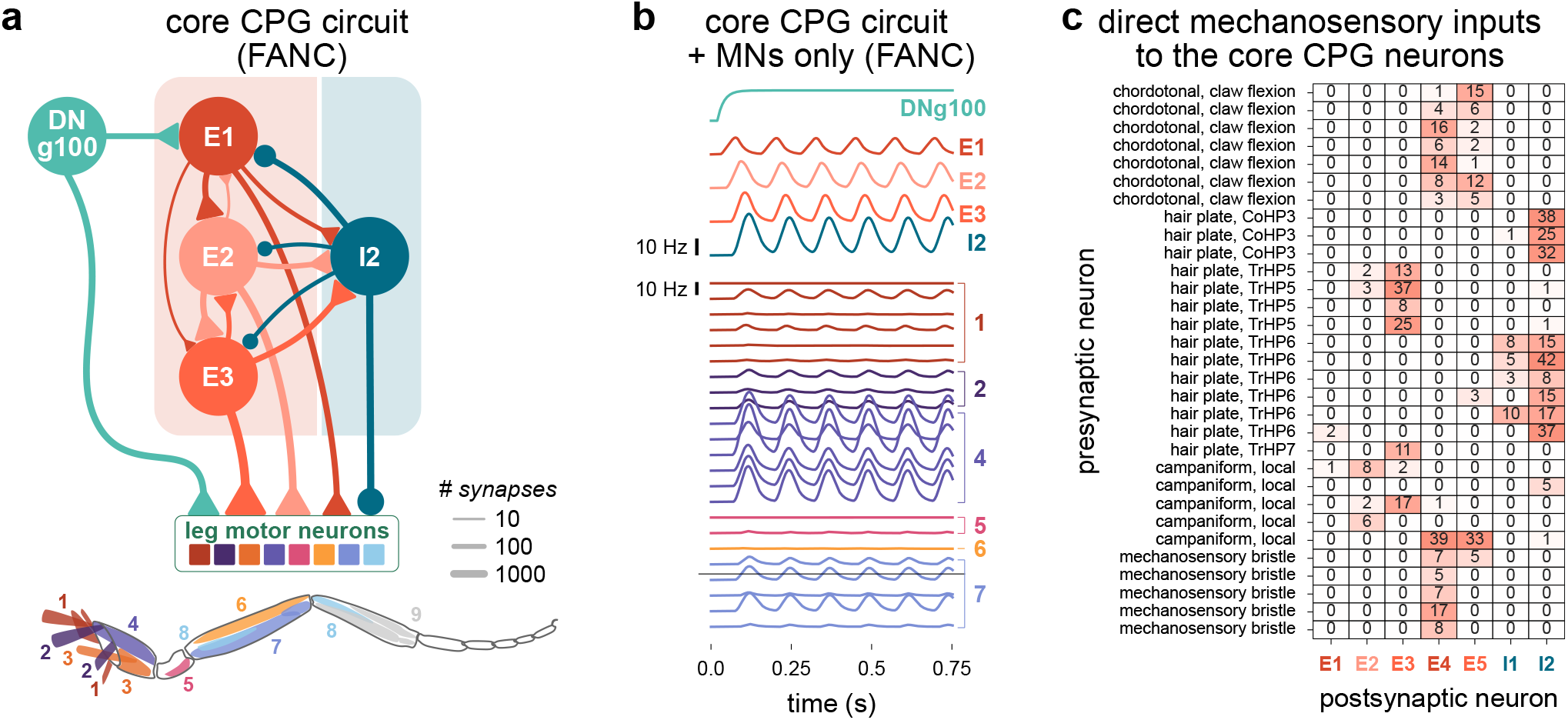
FANC core CPG circuit and mechanosensory inputs. **a**, The most common minimal circuit in FANC and its connectivity to motor neurons (shown are connections of at least 10 synapses). Triangles and circles are excitatory and inhibitory connections, respectively. Synapse counts are summed across all leg motor neurons. **b**, Example activity of the isolated four-neuron FANC core CPG circuit. **c**, All synapses from mechanosensory axons onto the core CPG neurons in FANC. Sensory neurons displayed are the complete set of sensory neuron axons belonging to either chordotonal organ neurons, hair plate neurons, campaniform sensilla neurons, or mechanosensory bristle neurons that make at least 1 synapse onto any of the *7* interneurons of interest.

**Extended Data Fig 8.**
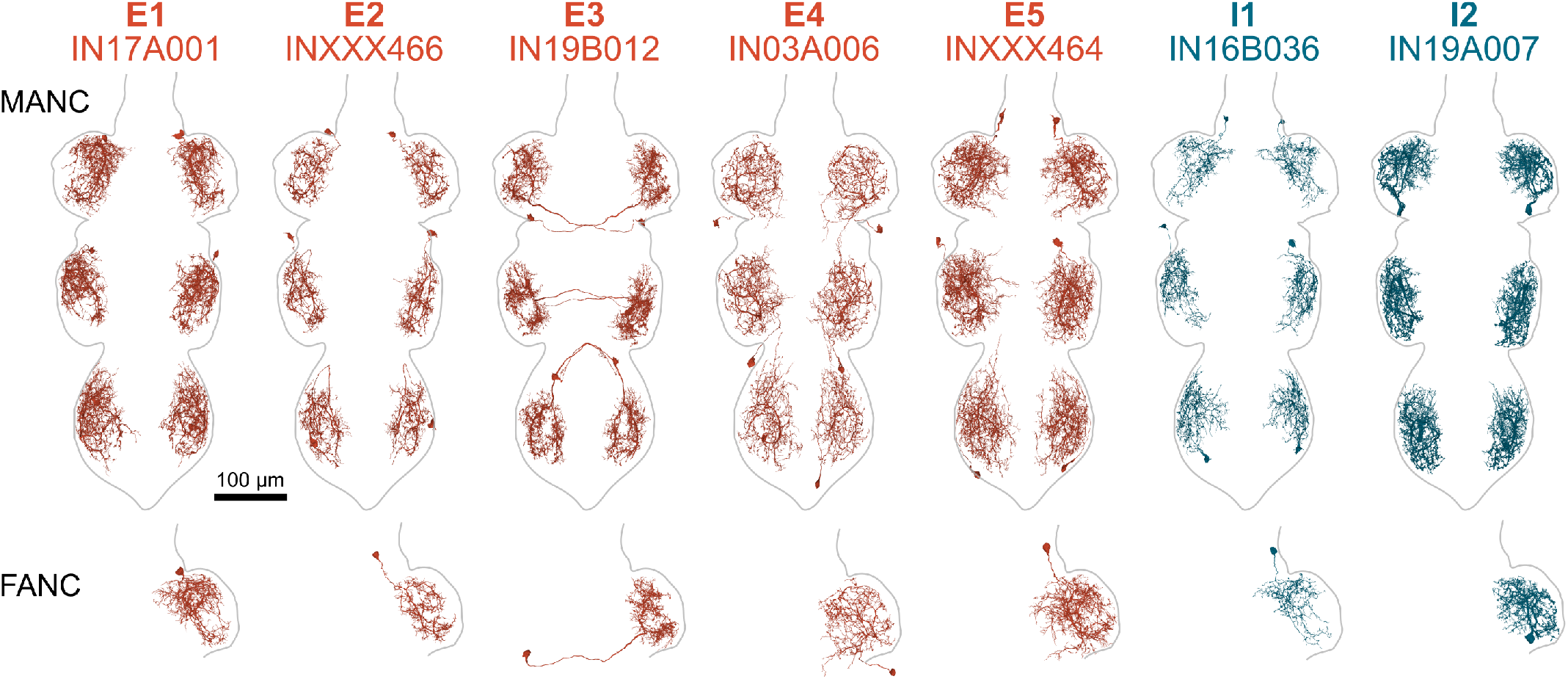
Morphologies of VNC interneurons identified in the computational sufficiency screens. Cell type correspond to the most commonly identified cells in **Supplementary Tables 3, 4, and 7**. *Top:* All neurons of each type in MANC. Each of these cell types consists of exactly one neuron per leg neuropil. *Bottom:* The neuron of the same type identified in the FANC front left leg neuropil.

**Extended Data Fig 9.**
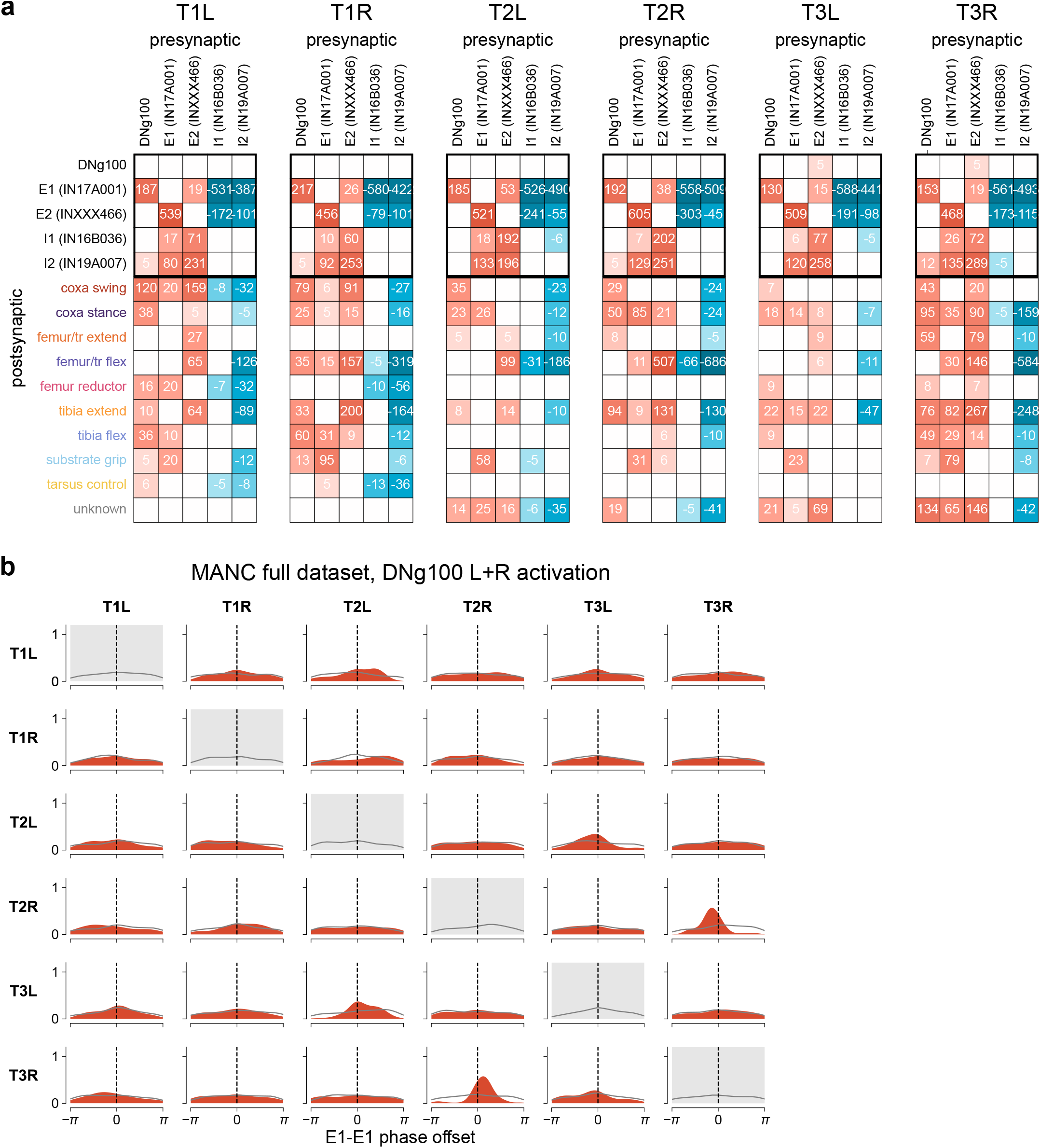
Connectivity and phase offsets of the core CPG circuit across six legs. **a**, Connectivity of the E1, E2, I1, and I2 neurons to each other and to the leg motor neurons in each of the six leg neuropils. Motor neurons have been grouped by module, with synapses counts summed within each module. Note that the number of motor neurons corresponding to each module changes in different legs. **b**, KDE plots showing the distribution of the six E1 neurons’ phase offsets relative to one another across the 128 replicates of bilateral DNg100 activation in the full MANC dataset (in particular, the offset of the E1 indicated by the column relative to the E1 indicated by the row). Gray lines are the distribution of a trial-shuffled control for each neuron pair.

**Extended Data Fig 10.**
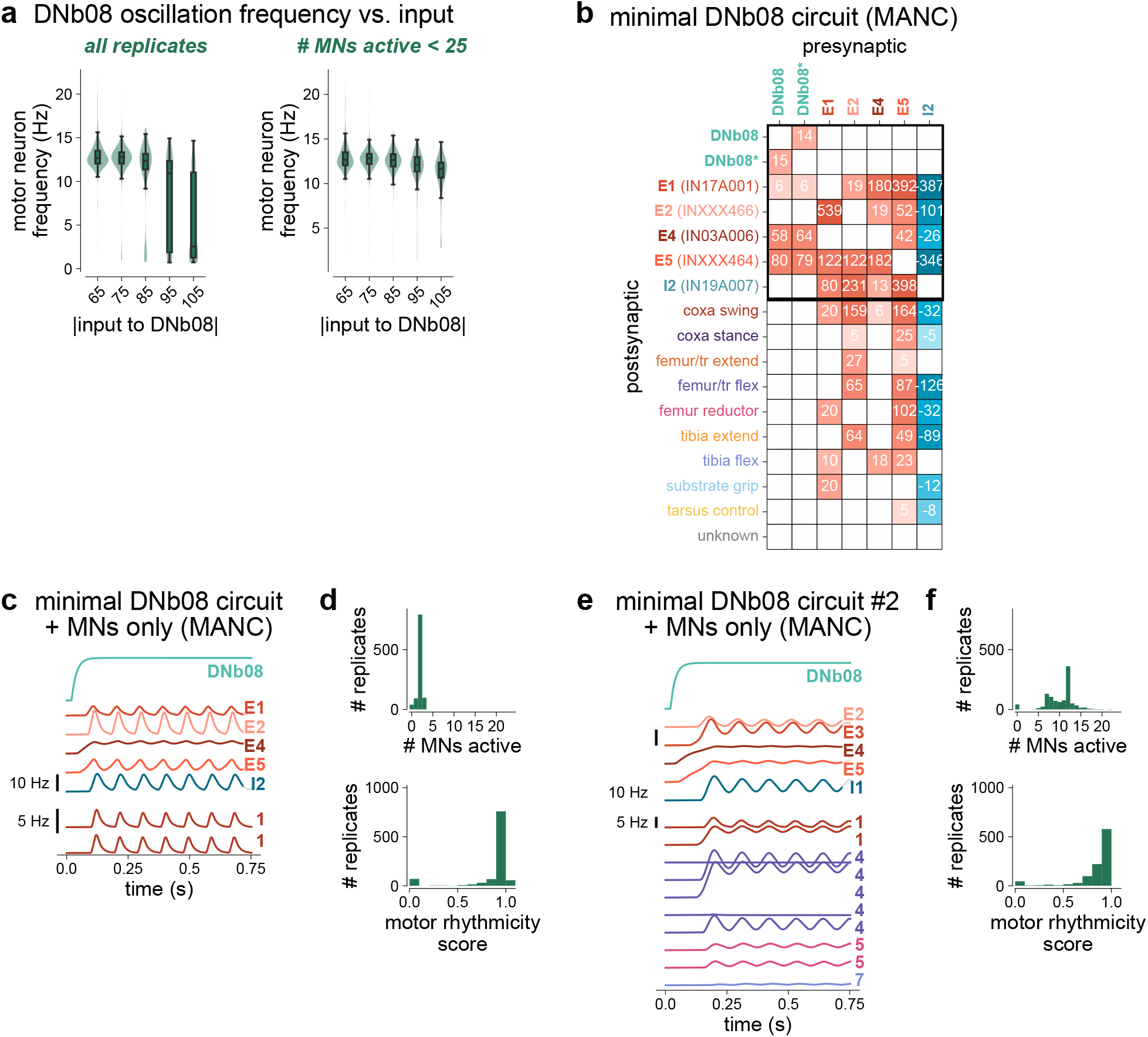
Additional analyses pertaining to DNb08. **a**, DNb08 activation does not show the modulation of motor neuron frequency with input strength (*n*=512 replicates per condition). Green shaded regions are distributions of frequencies, and black bars indicate medians and quartiles. Right plot excludes data in which *>*25 motor neurons were recruited, leaving *n*=512, 499, 435, 298, 176 replicates respectively for the 5 values of input strength. Despite removing highly active replicates, we did not observe an increase in frequency with increased input. **b**, Connectivity of the DNb08 minimal circuit neurons to each other and to the leg motor neurons innervating the front left leg. Motor neurons have been grouped by module, with synapses counts summed within each module. For reference, both DNb08 neurons innervating the left leg neuropil have been included; the one used in our simulations and analyses in **Figure 5** is marked with an asterisk. **c**, Example isolated activity of the most common minimal circuit downstream of DNb08. This activity looks fairly similar to the circuit downsstream of DNg100, and recruits the same coxa promotors. **d**, DNb08 activation consistently produced rhythms in this isolated circuit. **e–f**, Same as **c–d** but for the second most common circuit (**Supplementary Table 7**). This circuit also consistently drove rhythmic output and recruited more motor neurons.

## Notes

### Competing Interest Statement

The authors have declared no competing interest.

### Summary of Updates

We have made substantial revisions to the manuscript that fall under three different umbrellas: I. Robustness of simulation results across model parameters II. In-depth analyses of the structure of the putative core CPG circuit III. Replication of motor rhythms in 2 newly published fly CNS connectome datasets and in all legs IV: Accessibility of code repository

